# PAK-family kinases promote cell fusion irreversibility by preventing cell wall repair

**DOI:** 10.64898/2026.01.27.701985

**Authors:** Julia M Coronas-Serna, Sajjita Saha, Ana J Pérez-Berná, Sophie G Martin

## Abstract

Cell–cell fusion is critical for the sexual life cycle, as it drives the unidirectional transition between haploid and diploid phases. In fungi, fusion requires not only plasma membrane merging but also local cell wall removal. However, the integrity of the cell wall, which resists internal turgor pressure and is essential for survival, is normally monitored by a cell wall integrity pathway (CIP) involving Rho1 GTPase, protein kinase C (PKC) and MAPK signaling, which promotes damage repair. How the CIP allows localized cell wall degradation is not known. In the fission yeast *Schizosaccharomyces pombe*, we previously identified *pak2Δ* mutants that exhibit transient fusion: cells briefly fuse and exchange cytoplasmic contents, which signal catastrophic haploid meiosis, but then reseal their fusion pore. Here, we show that PAK-family kinase activity is essential for cell-cell fusion, with major and minor contribution for Pak2 and the essential Pak1, respectively. Strikingly, osmotic stabilization largely suppresses all PAK-associated fusion defects, pointing to a role in cell-wall remodeling. Pak2 accumulates at the fusion site, where it promotes fusion pore opening and expansion. In its absence, Rho1-GTP, PKC and beta-glucan synthase Bgs4 show elevated levels at the fusion site, and their partial loss-of-function rescues fusion. Pak1 mutants have elevated MAPK activity. Thus, Pak2 and Pak1 both antagonize the CIP. Furthermore, correlative light and cryo–soft-X-ray tomography reveals that resealed pores in PAK mutants rebuilt an intact cell wall. We conclude that PAKs enforce fusion directionality by antagonizing cell-wall repair mechanisms that otherwise restore separation between mating partners.

## Introduction

Cell-cell fusion drives the transition between haploid and diploid phases during eukaryotic sexual reproduction. By initiating the diploid phase, the opening of the fusion pore imposes directionality to the sexual life cycle. In turn, the zygote, or its progeny, reduces the genome back to the haploid state through meiosis. In the fission yeast *Schizosaccharomyces pombe*, we previously identified a mutant that exhibits transient cell fusion, reversing the normal directionality and leading to meiosis in haploid genomes ^1^. Wild-type haploid cells of P and M mating types fuse to form a diploid zygote that immediately forms stress-resistant haploid spores through meiosis ^2^. In contrast, a fraction of *pak2Δ* cell pairs transiently fuse but reseal their fusion pore. In the process, they exchange cytosolic information – leading to reconstitution of a transcription factor – that induces meiosis in haploid cells, with catastrophic consequences^1^. How does Pak2 promote the directionality of cell fusion?

Cell fusion necessitates not only plasma membrane merging, but also local removal of extracellular material. Fungal cells are encased in a protective cell wall that resists the internal turgor pressure ^3^. For cells to fuse, progressive wall thinning, as shown by ultrastructural and super-resolution imaging ^4,5^, relies on localized secretion of hydrolytic enzymes. The fusion focus, an aster-like array of linear actin filaments underlies the concentration of glucanase-containing vesicles to locally digest the cell wall at the point of contact, while maintaining overall cell integrity ^4,6–8^.

At all other times than cell fusion, cell wall integrity, monitored and repaired by the cell wall integrity pathway (CIP), is essential for fungal cell survival ^9,10^. Cell wall integrity is monitored by surface sensors, as illustrated by the clustering of Wsc1 upon compressive forces ^11^. These promote the activation of Rho-family GTPases, notably Rho1, which activate redundant Protein Kinases C (PKC) Pck1 and Pck2 ^12^. In turn, Rho1-GTP and PKC promote the activity of beta-glucan synthases for local repair, essential during growth, ^12–14^ and of a MAPK pathway for transcriptional response in response to stresses ^10,15,16^. Previous work in *S. cerevisiae* showed that the CIP antagonizes cell fusion ^17,18^, suggesting that repair may be locally inhibited to allow fusion.

Pak2/Shk2, which is induced upon stress or starvation ^19,20^, encodes one of two fission yeast p21-activated kinases (PAK) ^21,22^, conserved effectors of Cdc42 GTPase, which relieves auto-inhibition through CRIB domain binding ^21–26^. The other PAK, Pak1/Shk1/Orb2, is essential for viability and regulates cell polarity and division ^24,25,27–29^ by regulating Cdc42 ^30,31^ and phosphorylating regulators ^28,29,32–35^. The function of Pak2 is less understood, though it may share function and substrates with Pak1 ^20^, as Pak2 overexpression rescues *pak1Δ* cell viability ^21,22^. Cdc42 and PAKs have also been implicated in cell fusion in yeasts ^1,36,37^ and in eukaryotes without cell wall, for instance to promote myoblast fusion ^38^.

Here we ask how PAKs prevent fusion pore re-sealing. Both PAKs contribute to cell-cell fusion, with Pak2 playing the major role. Pak2 localizes to the fusion site, where its kinase activity promotes fusion pore opening and expansion, preventing re-sealing. Osmotic stabilization attenuated PAK mutant defects, suggesting cell wall regulation. Indeed, the Rho1-PKC pathway promoting cell wall synthesis is enhanced in *pak2Δ* and its downregulation rescues *pak2Δ* fusion defects. Pak1 also regulates the CIP, through a distinct mechanism. Furthermore, correlative light and cryo-soft-X-ray tomography (cryoSXT) reveals re-sealed PAK mutants with rebuilt cell wall at the cell-cell interface. Our data show that PAKs promote cell fusion directionality by suppressing cell wall repair, which otherwise reseals the pore.

## Results

### PAK-family kinase activity promotes cell-cell fusion

We previously showed that a fraction of pak2Δ cell pairs exhibit transient cytosolic connection, which reseals, leading to haploid meiosis in the P-cell partner ^1^. Indeed, at 30°C, 35% of *pak2Δ* cell pairs fail to fuse, and 7% undergo apparent transient fusion, leading to spore formation in one partner (Fig 1A-B). The remaining pairs successfully fuse, though with a delay (Fig S1A) ^1^, suggesting the existence of a partially redundant function.

**Figure 1.**
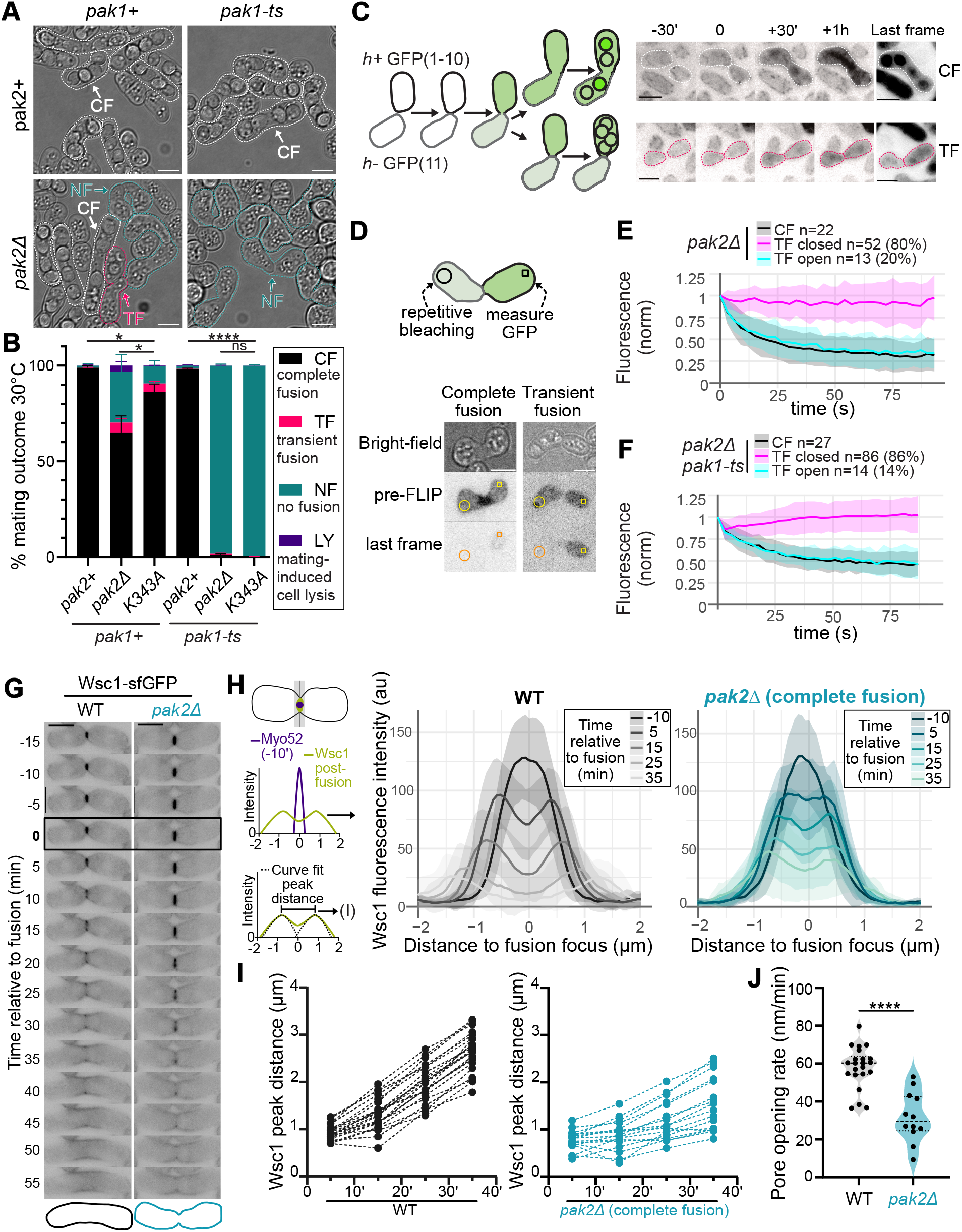
PAK-family kinases are required for cell fusion and pore expansion. (**A**) DIC images of mating outcomes at 30°C in wildtype, *pak2Δ*, *pak1-ts*, and double mutant: CF, complete fusion; TF, transient fusion; NF, no fusion. (**B**) Frequency of phenotypes in strains as in (A) and *pak2^K343A^* (kinase-dead); LY, mating-related lysis. Statistics: paired t-test on CF, with * p<0.05, **** p<0.0001 and ns p≥0.05. (**C**) Split-GFP to identify post-fusion pairs. Right: fluorescence timelapse of split-GFP in *pak2Δ* with complete (top) and transient (bottom) fusion. Last frame contrasted differently. (**D**) FLIP assay: cells were bleached in circles and measured in squares. Images of *pak2Δ*. (**E-F**) Normalized fluorescence intensity in *pak2Δ* (E) and *pak2Δ pak1-ts* (F) in CF, TF-closed and TF-open post-fusion cells. (**G**) Timelapse of Wsc1-sfGFP in wildtype and *pak2Δ* completely fused pairs, with zygote outline (55’). Fusion time is boxed. (**H**) Wsc1-sfGFP intensity across the fusion pore at indicated times; n=30 WT, n=21 *pak2Δ* CF pairs. Shaded areas show standard deviation. (**I**) Wsc1-sfGFP peak distance over time, measured from double Gaussian fitting on Myo52-centered curves. Dotted lines connect individual pairs; n=22 WT, n=17 *pak2Δ* CF pairs. (**J**) Pore opening rate from (I), WT n=22, *pak2Δ* n=12. Statistics: t-test with **** p<0.0001.

We probed the role of Pak1 during cell-cell fusion, using the temperature-sensitive allele *orb2-34* (named *pak1-ts* below), which exhibits reduced kinase activity even at permissive temperature ^30^. While *pak1-ts* cells exhibited round shapes, they fused as efficiently as wildtype cells (Fig 1A-B). Thus, shape and polarity defects do not per se compromise cell fusion.

By contrast, cell fusion was strongly compromised in *pak2Δ pak1-ts* double mutants at 30°C (Fig 1A-B), indicating that both PAKs redundantly contribute. A Pak2 kinase-dead allele (*pak2^K343A^*) showed a partial fusion defect, though weaker than that of *pak2Δ*, and *pak2^K343A^ pak1-ts* double mutants were as severely compromised as *pak2Δ pak1-ts* (Fig 1B). We conclude that PAK-family kinase activity is critical for cell fusion.

### PAKs promote fusion pore expansion post-fusion

Mutant cells that fuse but appear to reseal their pore suggest that PAKs also ensure irreversibility post-fusion. To test for complete re-closure, we developed a split-GFP system, with each partner cell expressing a single GFP fragment, leading to fluorescence only upon cytosolic mixing (Fig 1C). Thus, only pairs that fuse (transiently or not) are fluorescent.

Fluorescence loss in photobleaching (FLIP), bleaching one partner cell and recording fluorescence intensity in the other, led to rapid loss of fluorescence in completely fused pairs. By contrast, in over 80% GFP-positive *pak2Δ* and *pak2Δ pak1-ts* pairs with apparent cell wall at the cell-cell interface, no loss of fluorescence was observed, indicating full pore resealing (Fig 1D-F). Since fluorescence demonstrates cytosolic mixing and thus pore formation in both cell wall and plasma membrane, at least one of these openings is re-sealed.

Fusion pore expansion is likely limited by cell wall remodeling. We measured the speed of fusion pore opening with the cell wall sensor Wsc1-sfGFP, which accumulates at sites of cell wall compression and the fusion site ^11^. We observed similar levels of Wsc1 in *pak2Δ* and wildtype pairs before fusion (Fig 1G-H, Fig S1B), suggesting similar compressive forces. Post-fusion, Wsc1-sfGFP formed two peaks decorating the edges of the fusion pore (Fig 1G-H), whose distance increased linearly, with a rate of pore diameter increase of 58 ± 11nm/min (or ∼ 1µm /17 min) in wildtype cells (Fig 1I-J). This linear expansion occurred after an initial fast opening phase, consistent with near absence of very small pores in EM data ^5^. In fused *pak2Δ* cells, pore expansion was significantly slower, with a rate of 32 ± 9nm/min (or ∼ 1µm /31 min) (Fig 1H-J). Thus, Pak2 also functions after fusion pore opening to promote its expansion and prevent its re-closure.

### Pak2 activity is controlled at the fusion site to prevent cell lysis

Pak2 localizes to the fusion site (Fig 2A). Using the type V myosin Myo52-mScarlet-I as fusion focus marker ^7^ and detecting pore opening through exchange of P-cell-expressed cytosolic mTagBFP2, we measured Pak2-sfGFP fluorescence at the Myo52 position and synchronized traces at fusion time. Pak2 enrichment and localization were indistinguishable from Myo52 (Fig 2A-B), indicating co-localization with the actin fusion focus, like its activator Cdc42 ^37^.

**Figure 2.**
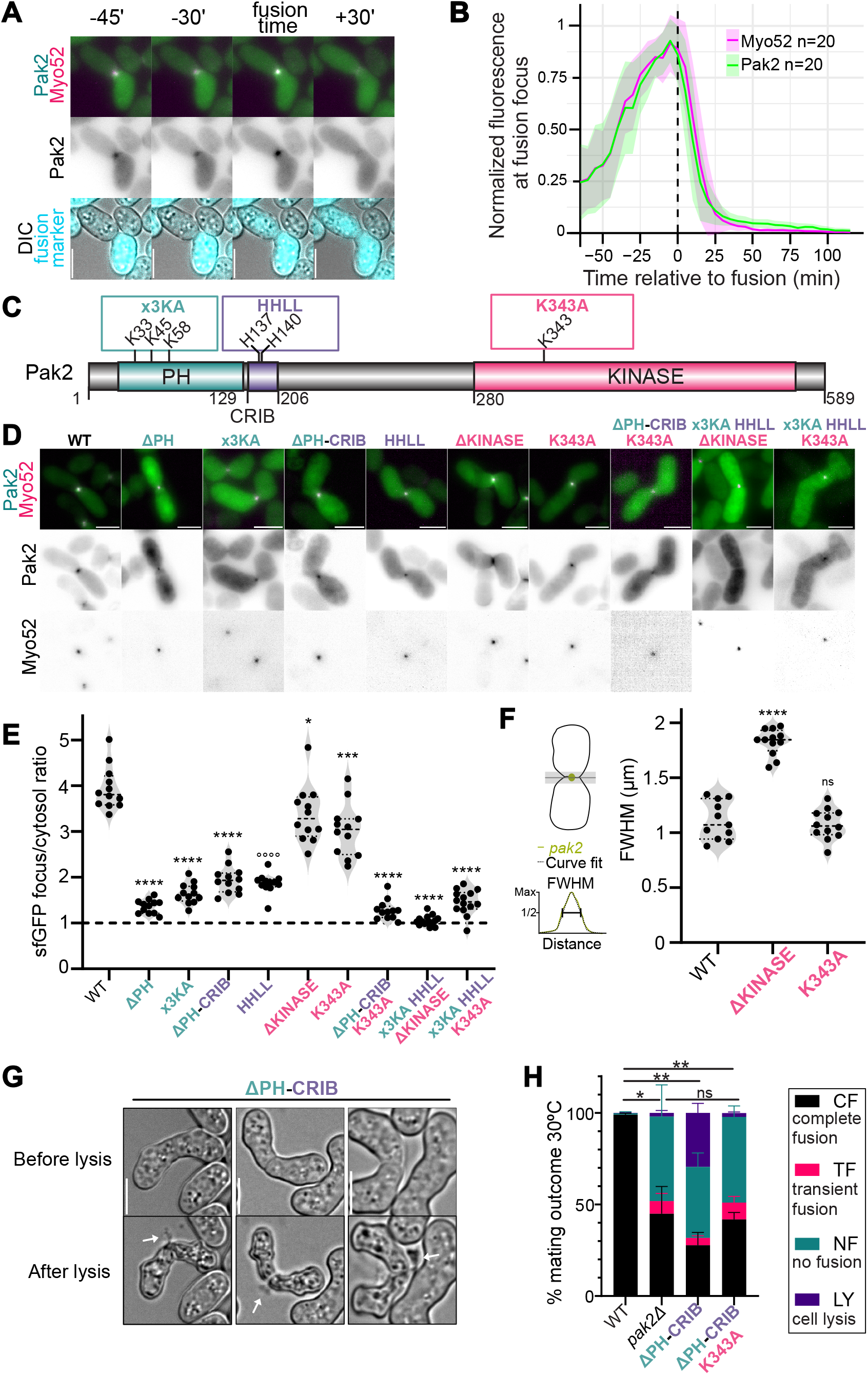
Pak2 activity is controlled at the fusion site to prevent cell lysis. (**A**) Timelapse of Pak2-sfGFP, Myo52-mScarlet-I, P-cell expressed mTagBFP2 and DIC during cell fusion. (**B**) Normalized fluorescence of Myo52-mScarlet-I and Pak2-sfGFP at the fusion focus (n=22). Shaded areas show standard deviation. (**C**) Pak2 protein domains and relevant residues. (**D**) Images of Pak2-sfGFP with indicated mutations and Myo52–mScarlet-I at t=-5’. (**E**) Pak2-sfGFP intensity at the fusion focus normalized to cytosol in cells as in (D). (**F**) Full width at half maximum (FWHM) of Pak2-sfGFP profiles across the fusion site. Statistics for (E-F): t-test (*) or Mann-Whitney test (°) vs WT with * p<0.05, *** p<0.001, ****/°°°° p<0.0001 and ns p≥0.05. (**G**) Lysis of *pak2^ΔPH-CRIB^*pairs, with cellular content extruded form the fusion site (arrows). (**H**) Frequency of mating outcomes at 30°C for indicated strains. Statistics: paired t-test on CF, with * p<0.05, ** p<0.005 and ns p≥0.05.

PAKs are autoinhibited by CRIB-mediated kinase suppression ^39–41^ (Fig 2C), until relieved by Cdc42-GTP binding ^42^. Pak2 further contains a N-terminal Pleckstrin Homology (PH) domain, whose deletion (pak2^ΔPH^), or mutation of its phospholipid-binding interface (pak2^x3KA^), reduced Pak2 enrichment at the fusion site (Fig 1C-E). Similarly, CRIB mutations preventing Cdc42-binding (pak2^HHLL^), or deletion of both domains (pak2^ΔPH-CRIB^), reduced Pak2 fusion site enrichment (Fig 2C-E). However, all mutants showed residual localization. Deletion of the kinase domain led to spreading of Pak2 over the cell-cell contact area, indicating a role in focalizing Pak2 (Fig 2C-F). In Pak2 mutants lacking PH and CRIB functionality, kinase domain deletion fully abrogated Pak2 localization (pak2^x3KA-HHLL-Δkinase^). However, Pak2 localization does not require kinase activity, as the kinase-dead K434A mutation neither led to Pak2 spreading, nor abrogated Pak2^ΔPH-CRIB^ or Pak2^x3KA-HHLL^ localization (Fig 2C-F). We verified that truncated Pak2 constructs are stable (Fig S1C), indicating that reduced localization is not due to protein degradation. In conclusion, the combined action of each domain, independently of kinase activity, drives Pak2 to the fusion site.

Deletion or mutation of any of Pak2 domains led to fusion defects (Fig S1D). Interestingly, while deletion of Pak2 PH domain showed defects similar to *pak2Δ* (Fig S1D), deletion of both PH and CRIB domains, predicted to relieve autoinhibition, showed a distinct phenotype of mating-associated lysis in 25% pairs, with cytosol extrusion from the cell-cell contact site (Fig 2G-H). The lysis phenotype was fully abrogated by the kinase-dead mutation K343A (Fig 2H). Our interpretation is that excessive local Pak2 activity at the fusion site leads to mating-associated lysis.

### PAKs does not strongly affect actin fusion focus formation

We probed whether Pak2 regulates the organization of the fusion focus. In all *pak2* mutant alleles, Myo52 localized at the fusion site as a dot, like in wildtype cells, reflecting the aster-like organization of actin filaments (Fig 2D). Similarly, Myo52 formed a dot in *pak2Δ* and strongly fusion-deficient *pak2Δ pak1-ts* mutants (Fig 3A). Thus, PAKs are not essential for the formation of the actin fusion focus.

**Figure 3.**
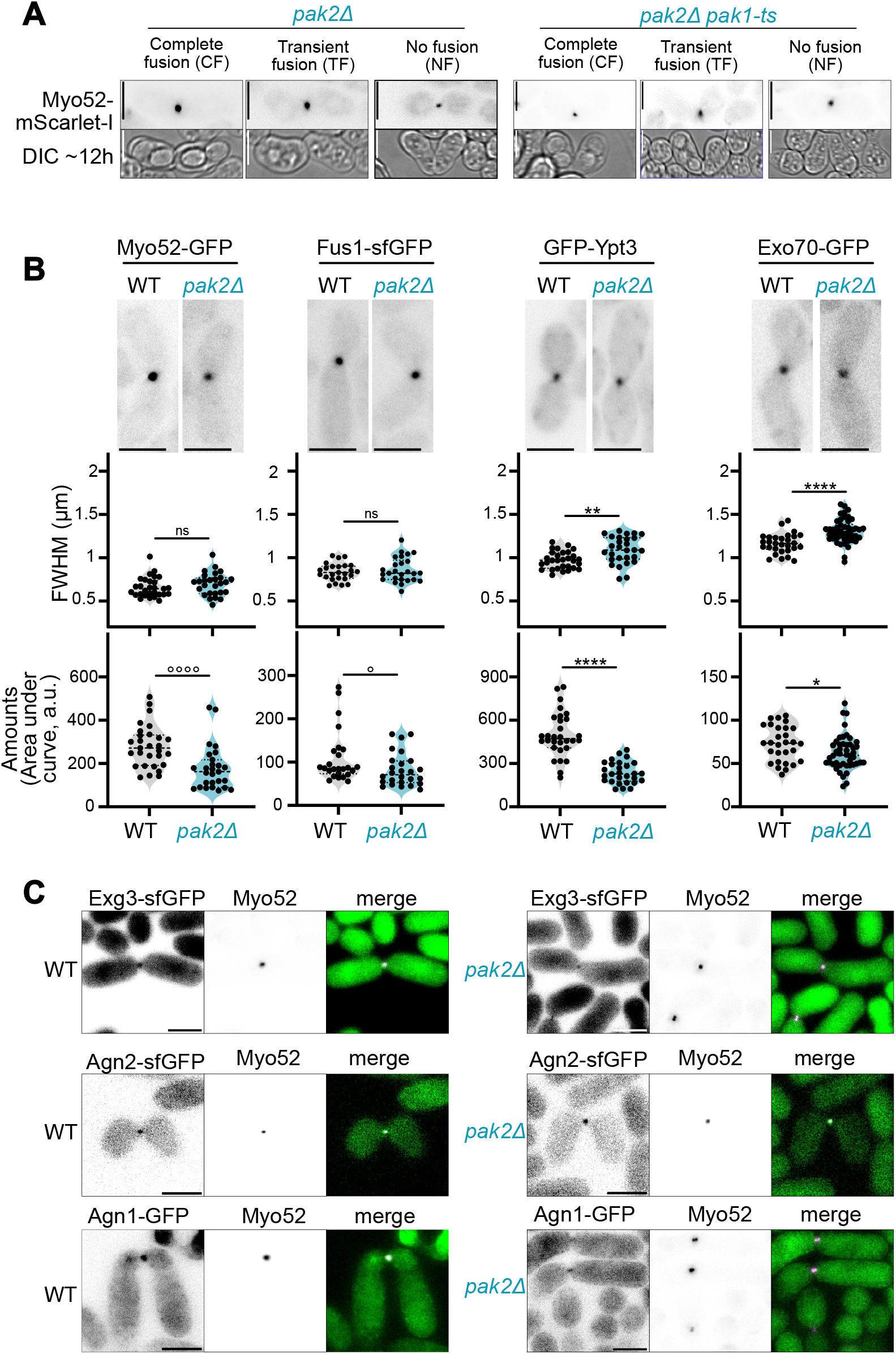
The actin fusion focus is not strongly altered in absence of PAKs. **(A)** Myo52-mScarlet-I (t=-10’) and DIC after ∼12h in *pak2Δ* and *pak2Δ pak1-ts* with indicated phenotype. No fusion pairs show a bright Myo52 frame. Images contrasted differently. (**B**) GFP-tagged Myo52, Fus1, Ypt3 and Exo70 in wildtype and *pak2Δ* pairs (t = -10’) (top), with corresponding FWHM (middle) and area under the curve (bottom) values. n≥25 each strain. Statistics: t-test (*) or Mann-Whitney test (°) with */° p<0.05, ** p<0.005 ****/°°°° p<0.0001 and ns p≥0.05. (C) Airyscan images of indicated GFP-tagged glucanases and Myo52-tdTomato in WT and *pak2Δ* pairs. Images contrasted differently.

To test if Pak2 plays a minor function, we quantified local fluorescence intensity and distribution width along the plasma membrane of Myo52 and the formin Fus1, which nucleates the fusion focus ^7^, 10min before fusion. Both proteins formed a similarly tight focus as in wildtype cells, though protein amounts were slightly reduced in *pak2Δ* cells (Fig 3B). The Rab GTPase Ypt3 and exocyst component Exo70, both of which decorate secretory vesicles ^4^, were also focused in *pak2Δ* cells, though with slightly more dispersed, lower intensity signal than in wildtype (Fig 3B). Finally, glucanases localized at the fusion site in both wildtype and *pak2Δ*, though signal intensity was too weak to quantify (Fig 3C). We conclude that *pak2Δ* cells show no gross abnormality in fusion focus organization and glucanase secretion. Previous work also showed that *pak2Δ* did not strongly affect fusion focus formation in the *cdc42-mCherry^SW^* mutant background, where cells are completely fusion-defective ^37^. These mild phenotypes are unlikely to fully explain the fusion defect and the transient fusion phenotypes of *pak2Δ* mutants.

### PAK mutant phenotypes are suppressed by sorbitol

Lysis in *pak2^ΔPH-CRIB^* cells may be a consequence of plasma membrane damage or cell wall integrity defect. Remarkably, sorbitol, which acts as osmo-stabilizer and alleviates cell wall damage-associated lysis, not only reduced lysis in *pak2^ΔPH-CRIB^* cells, but also strongly improved fusion success (Fig 4A, vs Fig 2H). Sorbitol similarly suppressed the fusion defects in *pak2Δ* and *pak2Δ pak1-ts* (Fig 4B). While wildtype cells fused slower in 1M sorbitol, *pak2Δ* did not slow fusion further in these conditions (Fig S1A). Thus, addition of an osmo-stabilizer largely alleviates the need for PAKs during cell fusion, suggesting that phenotypes in their absence are caused by cell wall problems.

**Figure 4.**
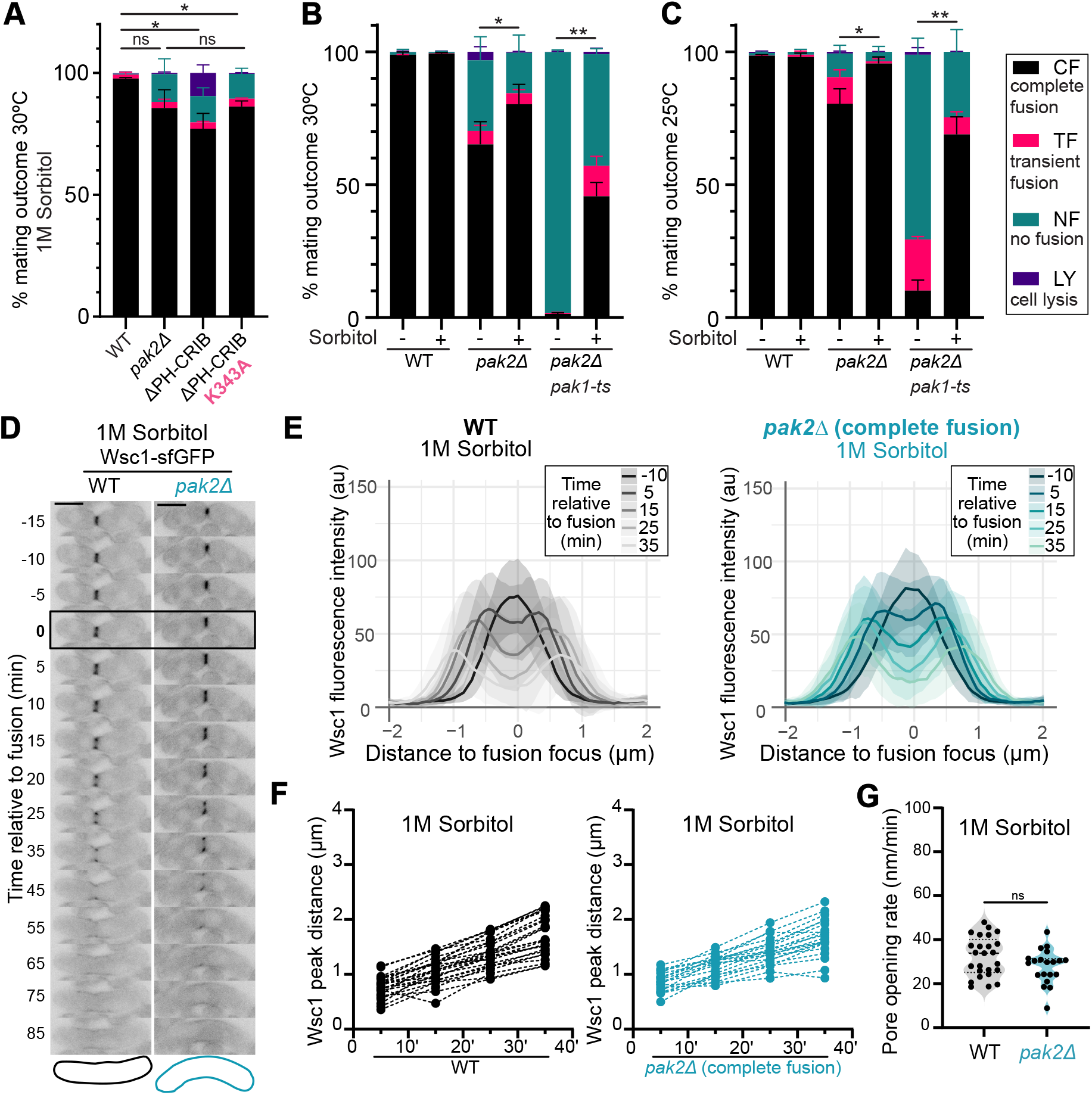
Suppression of PAK mutant phenotypes by sorbitol. (**A-C**) Frequency of mating outcomes of indicated strains at 30°C (A, B) or 25°C (C) with or without 1M sorbitol. Statistics: paired t-test on CF, with * p<0.05, ** p<0.005, ns p≥0.05. (**D**) Timelapse of Wsc1-sfGFP on 1M sorbitol in WT and *pak2Δ* completely fused cell pairs, with zygote outline (85’). Fusion time is boxed. (**E**) Wsc1-sfGFP intensity across the fusion pore at indicated times, with 1M Sorbitol, quantified as in Fig 1H; n=31 WT, n=24 *pak2Δ* CF pairs. Shaded areas show standard deviation. (**F**) Wsc1-sfGFP peak distance over time with 1M sorbitol, measured as in Fig 1I. Dotted lines connect individual pairs; n=26 WT, n=21 *pak2Δ* CF pairs. (**G**) Pore opening rate with 1M sorbitol from (F), WT n=25, *pak2Δ* n=20. Statistics: t-test with ns p>0.05.

In parallel experiments, we discovered that PAK fusion phenotypes are temperature dependent. Mating at 25°C reduced *pak2Δ* fusion failures to only 10%, with another 10% undergoing transient fusion ^1^ (Fig 4C, S1E). Similarly, a small fraction of *pak2Δ pak1-ts* double mutants successfully fused, and 20% showed transient fusion at 25°C (Fig 4C). Addition of sorbitol suppressed these defects at both 25°C and 30°C (Fig 4B-C). Conversely, temperature increase to 32°C caused >90% fusion failure in *pak2Δ* (Fig S1E). Thus, temperature and sorbitol have opposite influence on the PAK function, respectively compounding and relieving the fusion defect.

We further tested the influence of sorbitol on pore expansion. In wildtype cells, sorbitol slowed pore expansion rates to 32 ± 13 nm/min, suggesting that high turgor promotes pore expansion. Remarkably, *pak2Δ* cells showed similar rates (28 ± 8 nm/min; Fig 4D-G). Thus, Pak2 activity is required to promote the high pore-opening rates that occur in normal conditions (i.e. without sorbitol). In contrast, sorbitol slows both fusion and pore expansion (Fig 4G, S1A) and alleviates the requirement for PAK activity.

The temperature and sorbitol sensitivities of PAK mutants, especially *pak2*Δ, suggest these environmental changes alter the cellular requirement for these kinases. Both conditions affect the cell wall. Notably, high temperature promotes CIP activation ^10,16,43,44^, whereas sorbitol provides osmotic support that rescues the growth of cells with reduced cell wall biogenesis ^45–50^. Sorbitol also reduces mechanical forces on the cell wall ^51^, likely reducing activation of the CIP^11^. Indeed, Wsc1 accumulation at the pre-fusion site was reduced in both wildtype and *pak2Δ* grown on sorbitol (Fig S1B). These observations strongly suggest that PAKs promote cell fusion by inhibiting cell wall biogenesis and/or repair.

### PAK mutants exhibit increased CIP activity

We tested whether Pak2 regulates the CIP. To probe Rho1 activity, we designed a Rho1-GTP biosensor by expressing the Pck2 Rho-binding domain (RBD) fused with sfGFP (Fig S2A), named RBD2-sfGFP, similar to previously built sensors ^52,53^. In mitotic cells, at restrictive 36°C, RBD2-sfGFP localized at division site and cell poles in wildtype but not *rho1-596* temperature-sensitive (*rho1-ts* below) cells, in which Rho1-GTP levels are strongly reduced ^50^ (Fig S2B). RBD2-sfGFP signal was also reduced at 25°C (Fig S2C), consistent with *rho1-ts* being hypomorphic at permissive temperature ^50^. Localization was not affected by deletion of *rho2*, nor either of *rho3*, *rho4* or *rho5* (Fig S2C-D). During cell fusion, RBD2-sfGFP localized to the cell-cell contact site, with local intensity at the Myo52-labelled focus increase until peak at fusion time and drop as the fusion pore expanded (Fig 5A-B). In *pak2Δ*, RBD2-sfGFP intensity was higher at the fusion site throughout (Fig 5A-B), but unchanged at the division site (Fig S2E). A different quantification method confirmed elevated levels of Rho1-GTP in *pak2Δ* pre-fusion cells, irrespective of mating outcome (Fig 5C-D, 5F). As Wsc1 sensor levels are unchanged (see Fig 1H, Fig S1B), this indicates that Pak2 downregulates Rho1 activation downstream of Wsc1. In post-fusion cells, RBD2-sfGFP signal remained high in *pak2Δ*, likely reflecting slow pore expansion or re-closure (Fig 5E-F). We conclude that Pak2 antagonizes Rho1-GTP activation at the fusion site.

**Figure 5.**
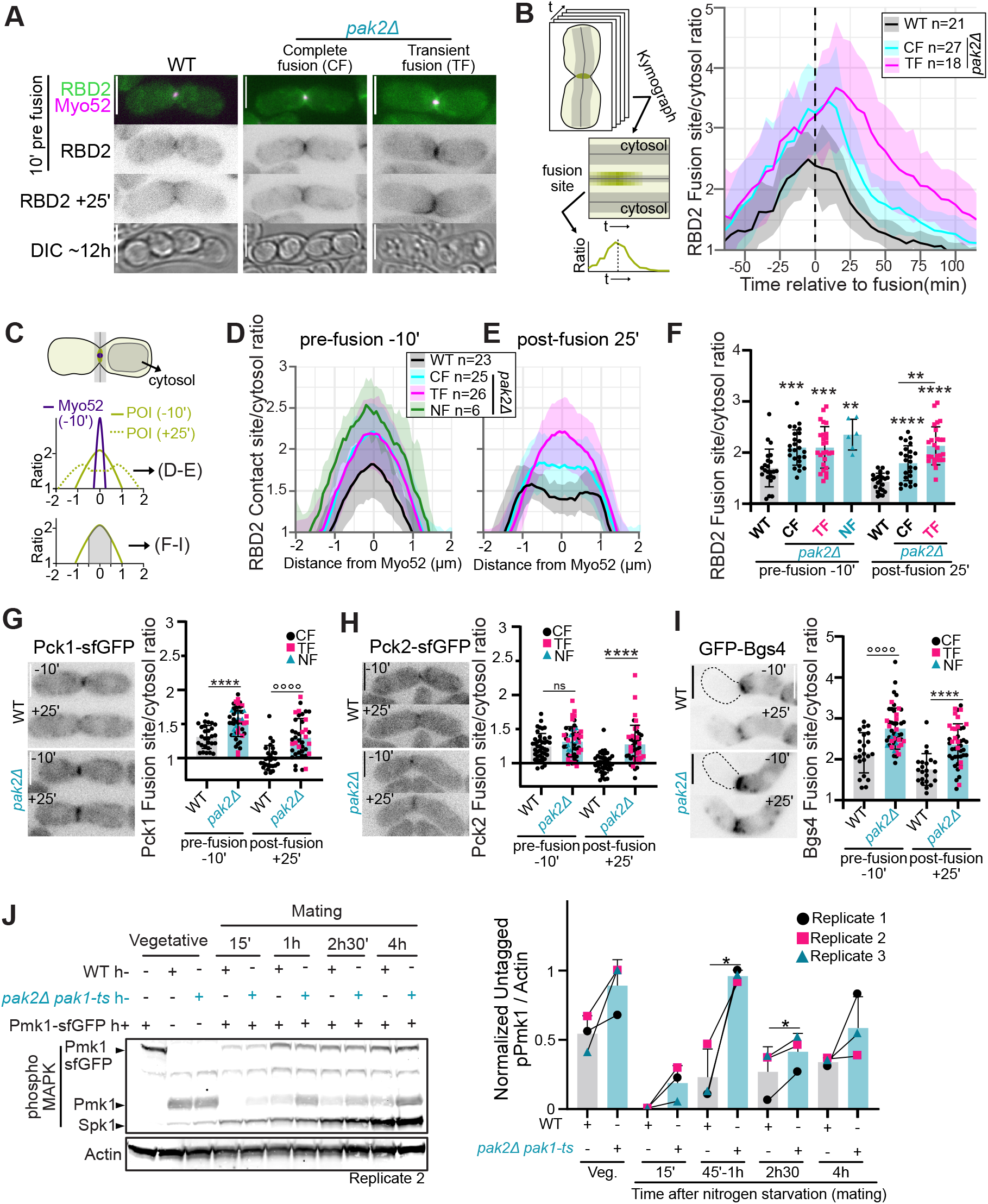
Hyper-activation of the CIP in PAK mutants. **(A)** Images of RBD2-sfGFP and Myo52-mScarlet-I at indicated timepoints and DIC after ∼12h, in WT and *pak2Δ* with indicated phenotypes. (**B**) RBD2 at the fusion site over time, measured at the Myo52-marked focus from kymographs (left). Shaded areas show standard deviation. (**C**) Measurement of proteins in (D-I). Profiles at the contact site, aligned to the Myo52 focus and divided by cytosolic signal are shown in (D-E). Mean of the fusion site (medial 1µm) values are shown in (F-I). (**D**-**E**) RBD2 contact site to cytosol ratio, pre- (D) and post-fusion (E). For No Fusion (NF) pairs, a bright Myo52 frame was selected. Shaded areas show standard deviation. (**F**) RBD2 fusion site to cytosol ratio, in WT and *pak2Δ* pairs, for indicated timepoints and phenotypes. (**G-I**) Images of Pck1-sfGFP (G), Pck2-sfGFP (H) *h90* cells and GFP-Bgs4 *h+* cells crossed to untagged *h-* (I), in WT or *pak2Δ* cells (left) and corresponding fusion site to cytosol ratio, pre- and post-fusion (right). Statistics: t-test (*) or Mann-Whitney test (°) with * p<0.05, ** p<0.005, *** p<0.001 ****/°°°° p<0.0001 and ns p≥ 0.05. (**J**) Phospho-MAPK western blot of untagged *h-* WT, *pak2Δ pak1-ts* and *h+* WT Pmk1-sfGFP strains grown individually or mated together for indicated time. Note the higher levels of phospho-Pmk1 in *pak2Δ pak1-ts* mutants, but equal levels in the WT partner. Phospho-Spk1 increase shows mating induction ^82^. Actin serves as loading control (bottom). Quantification on the right, with error bars for standard deviation. Statistics: paired t-test with * p<0.05.

Downstream of Rho1, both PKCs localized to the fusion site (Fig 5G-H). Pck1, like the RBD2 biosensor, showed elevated levels in *pak2Δ* before and after fusion (Fig 5G). Pck2 was not significantly different in *pak2Δ* and wildtype pre-fusion (Fig 5H). Both Rho1 and PKC promote beta-glucan synthase activity: beta-glucan synthase Bgs4 levels were also increased at *pak2Δ* fusion sites (Fig 5I). By contrast, phosphorylation of the downstream MAPK^Pmk1^, which marks kinase activation, appeared unchanged in *pak2Δ* cells (Fig S2F), perhaps because transient signaling increase at fusion sites is averaged out in extracts from unsynchronized cells. We conclude that Pak2 acts locally at the plasma membrane to negatively regulate CIP upstream factors.

In contrast to *pak2Δ*, *pak1-ts* did not cause RBD2-sfGFP increase, even in strongly fusion-deficient *pak2Δ pak1-ts* (Fig S2G). However, Pak1 regulated phospho-MAPK^Pmk1^ levels. Nitrogen starvation causes an acute reduction in phospho-MAPK^Pmk1^, which slowly increased again over time. In *pak1-ts* and *pak2Δ pak1-ts* mutants, phospho-MAPK^Pmk1^ remained higher and increased back to higher levels than in wildtype, in cells starved with or without mating partners (Fig 5J, S2H-I). Phospho-MAPK^Pmk1^ was not significantly affected by PAKs during mitotic growth, as previously reported ^16^. Thus, Pak1 negatively regulates the CIP during starvation but differently from Pak2.

In summary, PAK inactivation promotes CIP activation, setting PAKs as negative regulators of this pathway during sexual reproduction.

### Inactivation of the upstream CIP components suppresses *pak2Δ* fusion defects

To probe whether CIP hyper-activation causes the *pak2Δ* fusion defects, we tested whether its partial inactivation would restore fusion ability. Because *rho1* is an essential gene, we used *rho1-ts*, which reduces Rho1-GTP levels even at permissive temperature ^50^ (Fig S2C). At semi-permissive temperature, though a large fraction of cells lysed, *rho1-ts* significantly rescued *pak2Δ* fusion ability at 32°C and more modestly at 30°C (Fig 6A). Deletion of *pck1* or *pck2*, though individually viable, leads to cell lysis ^54^. Likely for this reason, we were unable to obtain a *pck2Δ pak2Δ* double mutant, but the *pck1Δ pak2Δ* double mutant showed improved fusion relative to *pak2Δ* (Fig 6B). Finally, four hypomorphic alleles in the essential beta-glucan synthase Bgs4 ^55–59^ restored fusion capacity to *pak2Δ* cells, to various extent (Fig 6C). We conclude that Pak2 promotes cell fusion by downregulating cell wall biosynthesis.

**Figure 6.**
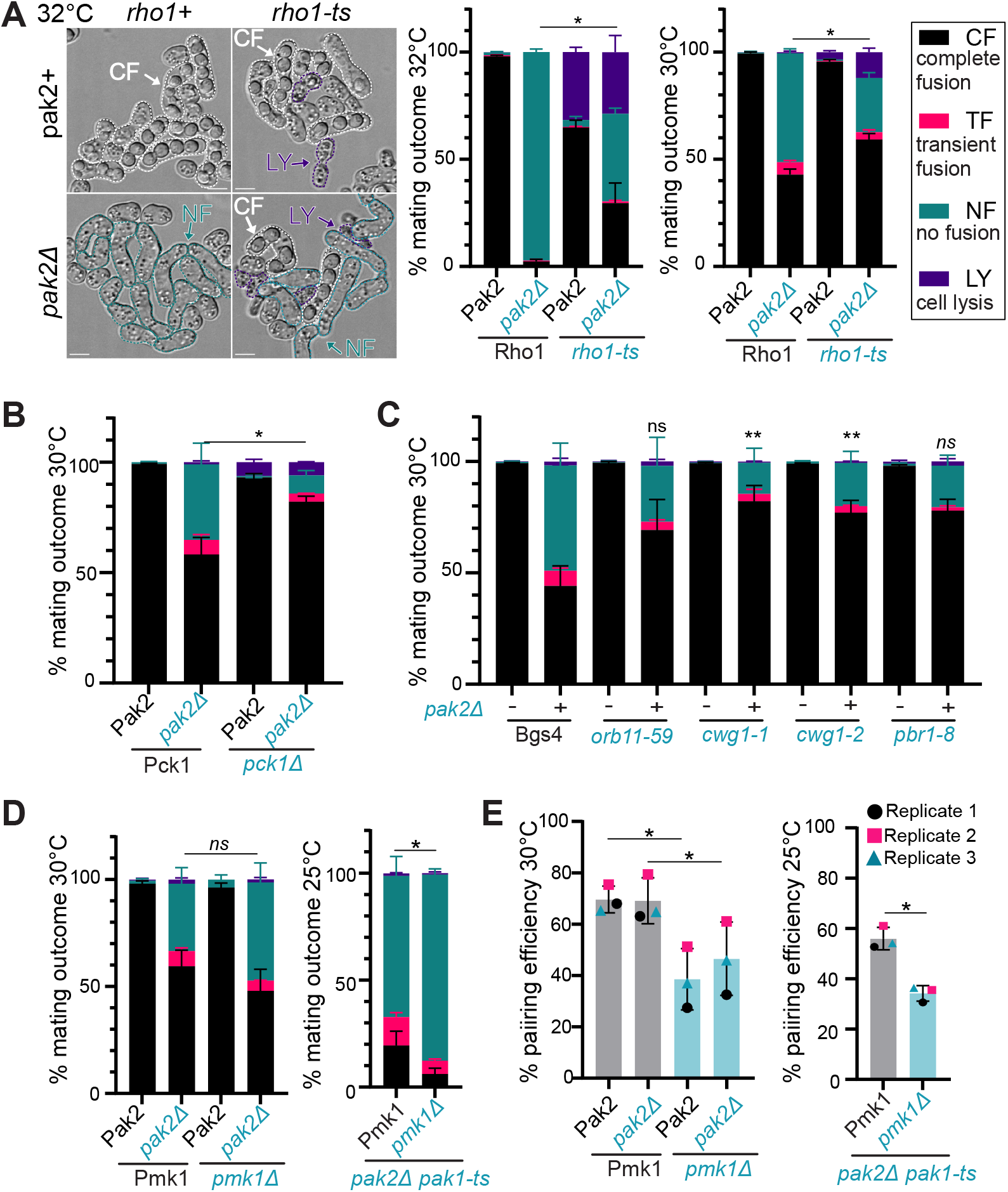
Suppression of *pak2Δ* fusion phenotype by CIP mutants. (**A-D**) Images and frequency of mating outcomes at the indicated temperatures and strains, comparing the effect of *rho1-ts* (A), *pck1Δ* (B), *bgs4* hypomorphic alleles (C) and *pmk1Δ* (D) in otherwise WT, *pak2Δ,* and *pak2Δ pak1-ts* backgrounds. Statistics: paired t-test or Wilcoxon test (italicized) on CF compared to *pak2Δ* or *pak2Δ pak1-ts*, with * p<0.05, ** p<0.005, ns/*ns* p≥0.05. (**E**) Cell pairing efficiency at the indicated temperature and strains. Statistics: paired t-test with * p<0.05.

Deletion of the MAPK Pmk1 did not rescue fusion in *pak2Δ* mutants (Fig 6D); *pmk1Δ* also did not improve fusion in *pak2Δ pak1-ts* double mutants. We note that *pmk1Δ* reduced the fraction of paired cells, suggesting crosstalk with the pheromone-MAPK pathway (Fig 6E). Nevertheless, these data suggest that hyper-activation of the CIP MAPK, while reflecting pathway dysregulation in PAK mutants, is not the cause of their fusion defect.

### Cryo-soft X-ray tomography reveals intact cell wall in re-sealed post-fusion PAK mutants

To directly observe cell wall repair in transiently fused cell pairs, we imaged full-volumes of fusion pairs by cryo-soft-X-ray tomography (cryo-SXT) (Fig 7A). Compared to electrons, X-rays penetrate deeper into the sample. This allows to reconstruct an image of the full cell volume from tomography without sectioning, with a resolution of ∼30 nm ^60^. Cryo-SXT of wildtype zygotes revealed clear ultrastructure, with easily identifiable fused nuclei, vacuoles, lipid droplets, cell wall, and a wide fusion pore (1.46 µm ± 0.52, n =7) (Fig 7B, Movie S1, Fig S3A). To identify transiently fused PAK mutant pairs, we used the split-GFP system, where cytosolic mixing reconstitutes fluorescence. Transient fusion often leads to asymmetric nuclear GFP fluorescence in the two partner cells, providing further identification of transiently fused pairs (Fig 1C). M-cells also expressed Myo52–mScarlet-I, labelling fusion focus and mating types (Fig 7A). We analyzed *pak2Δ pak1-ts* double mutant cells at 25°C, in which 60% of fused pairs re-seal their pore (Fig 4C).

**Figure 7.**
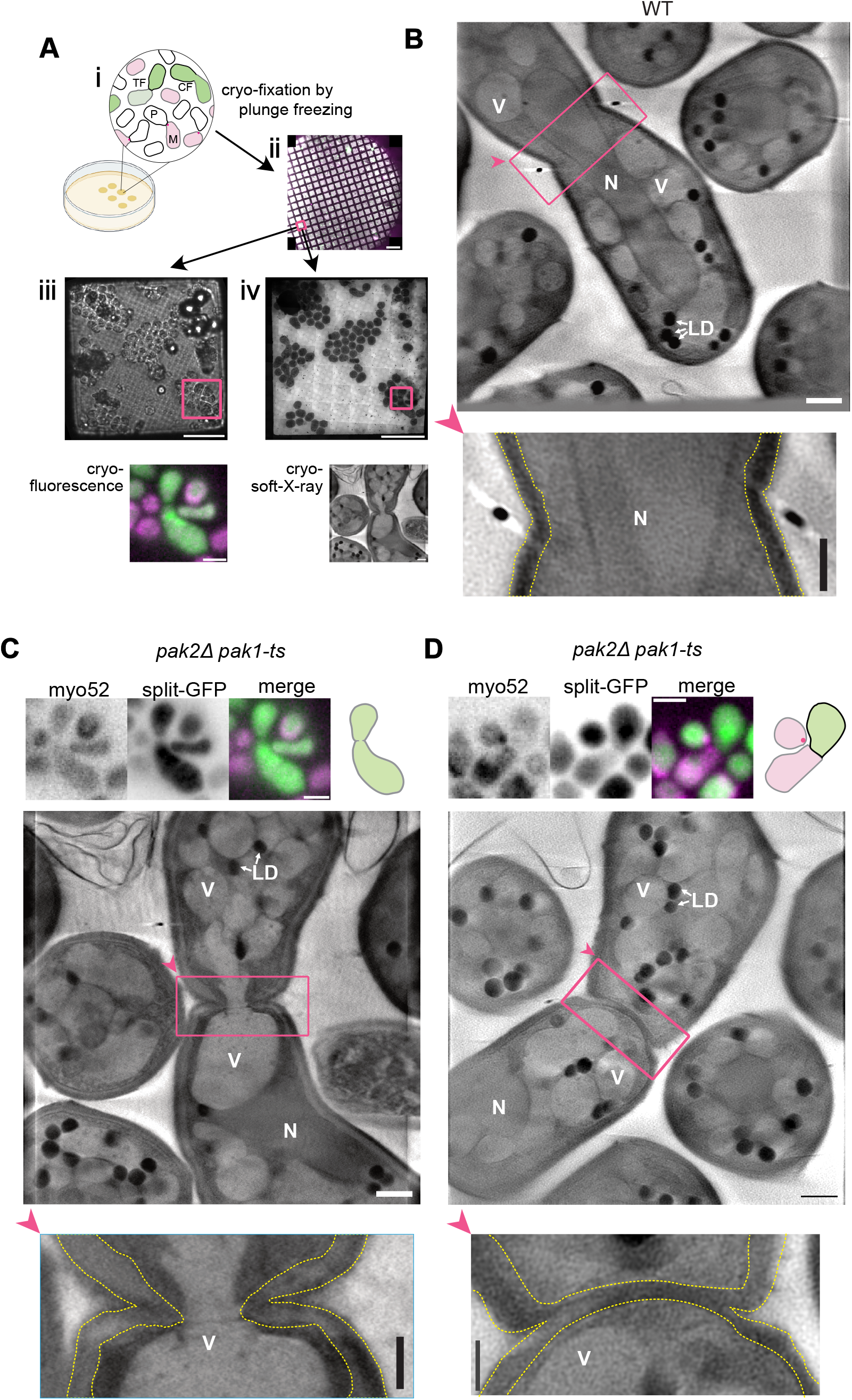
Complete cell wall re-building in post-fusion PAK mutants. (**A**) Correlative-light cryo-soft-X-ray tomography (cryo-SXT) sample preparation. (i) A Split-GFP tagged mating mixture was plated on N-starvation conditions (25°C), then resuspended and cryo-fixed by plunge freezing on grids (ii). Upon cryo-fluorescence microscopy imaging, GFP positive pairs were annotated (iii, scale bar 25 µm, enlarged square 5 µm). At the MISTRAL beamline at ALBA synchrotron, selected pairs were identified and targeted for cryo-SXT (iv, scale bar 25 µm, enlarged square 1 µm). (**B**) Cryo-SXT tomogram virtual slice of post-fusion WT pair, with expanded pore and fused nuclei. (**C-D**) Correlative light-cryo-SXT tomograph virtual slices of post-fusion *pak2Δ pak1-ts* pairs with a narrow, opened pore plugged by a vacuole (C), or a transiently fused and resealed pore (D). Top images show fluorescence and schematics. Arrowheads indicate the enlarged region orientation. Scale bars of fluorescence panels = 5 µm, full cryo-SXT = 1 µm, and enlarged region = 0.5 µm. N = nucleus, V = vacuole, LD = lipid droplets. The cell wall is indicated with yellow dotted lines.

Cryo-SXT full volume of 14 GFP-positive *pak2Δ pak1-ts* zygotes, which had fused in their past, revealed two classes of phenotypes. 7 pairs had an open fusion pore, smaller than that of wildtype (0.59 µm ± 0.17, n =7) (Fig S3A), consistent with slow pore expansion. These narrow fusion pores were plugged by a large organelle, either the nucleus or a vacuole (Fig 7C, Movie S2, Fig S3B). Remarkably, the other 7 pairs had no open pore and an apparently intact cell wall across the whole surface between partner cells (Fig 7D, Movie S3, Fig S3C). Thus, despite having opened the cell wall and merged their plasma membrane in the past, these pairs not only underwent membrane fission, closing the fusion pore, but also rebuilt the cell wall. We conclude that re-sealing of the fusion pore is driven by cell wall repair in absence of PAK regulation.

## Discussion

We show here that PAKs in fission yeast prevent cell wall biogenesis and repair at the fusion site. When their function is reduced, cell fusion either fails or occurs with slow pore expansion or its rapid resealing. These heterogeneous phenotypes can all be explained by overactive cell wall repair, which antagonizes local digestion to open the fusion pore or expand it post-fusion. Several lines of experimental evidence support this view. First, the proportion of each phenotype is strongly shifted by temperature and osmo-stabilization, two conditions that modulate the CIP. Second, the CIP is overactive in PAK mutant cells, as illustrated by elevated Rho1-GTP, PKC and Bgs4 at *pak2Δ* fusion sites and increased phospho-MAPK^Pmk1^ in *pak1-ts*. Third, downregulation of CIP components substantially rescues *pak2Δ* cell fusion. Finally, cryoSXT demonstrates cell wall rebuilding after a pore was created. Thus, PAKs inhibit the CIP to prevent cell wall repair and promote irreversible cell fusion.

### PAKs inhibit cell wall repair at the fusion site

The cell wall is essential for fungal cell viability. To ensure its integrity, the CIP monitors it through surface sensors and promotes its repair through recruitment of synthases ^10,11^. This activity is critical to repair acute damage ^61^, and maintains homeostatic wall thickness at sites of growth ^62^. During cell fusion, wall thinning requires the local concentration of hydrolytic activity, through secretory vesicle capture on the fusion focus and local secretion of their glucanase cargoes ^4,7^. Our data now show that wall thinning – and maintenance of an open fusion pore – also requires downregulation of cell wall biosynthetic activity by PAKs. In absence of PAKs, excessive CIP activity prevents local wall piercing and/or reverses the process leading to re-closure of the fusion pore. Thus, the CIP antagonizes cell fusion in fission yeast, as in budding yeast ^17,18^. We conclude that cell fusion not only requires local wall digestion but also downregulation of repair pathways.

CIP activation at the fusion site – recruitment of Wsc1, Rho1-GTP, PKC, Bgs4 – resembles that observed at damage sites ^61^, raising the question of how the cell distinguishes the fusion site from an injury, to repair only the latter. The cell-cell contact site differs from other surface regions in at least two ways. First, partner cells physically touch, such that repair downregulation, likely through PAKs, may be a consequence of physical engagement between surface factors. Recent work in *S. cerevisiae* has shown that the adhesin Fig2 serves as contact sensor to prevent PKC localization to contact sites ^63^, lending support to this hypothesis. A second difference is pheromone signaling, which occurs locally at the fusion site and promotes fusion ^64–66^. Pheromone signaling may not only promote Pak2 expression ^19^ but also local function to downregulate repair. In either case, PAKs ensure that cell wall digestion at the fusion site is not interpreted as an injury.

### Redundant, yet distinct roles of PAKs on the CIP

Although PAKs are redundantly required to allow fusion, how they regulate the CIP appears distinct. Pak2 localizes and acts at the fusion site: an allele lacking autoinhibition induces lysis at the fusion site; and *pak2Δ* causes local increase in Rho1-GTP, PKC and Bgs4 levels, which in turn prevent cell fusion. As Wsc1 accumulates in response to compressive forces ^11^ and its levels at fusion sites are not altered by Pak2, this indicates that Pak2 does not alter the force that engages the pathway but acts downstream. A link between Pak2 and the CIP was made previously, as expression of Pak2 kinase domain in mitotic cells caused lethality, which was suppressed CIP MAP3K^Mkh1^ deletion ^67^. However, further work found no evidence of MAPK^Pmk1^ activity change in *pak2Δ* ^16^, as we also observed during mating. We propose that Pak2 acts locally on upstream CIP components, for instance by regulating Rho1 GEFs and/or GAPs.

The small changes in vesicle distribution suggests that Pak2 may also slightly reduce glucanase secretion, contributing to the *pak2Δ* fusion defect. However, the increased levels of Bgs4 at the fusion site, itself a vesicular cargo ^55^, indicate that polarized secretion is not strongly perturbed. The effect of Pak2 on the actin fusion focus is small, as this structure forms largely correctly in fusion-defective *pak2Δ pak1-ts* and *pak2Δ cdc42-mCherry^SW^* double mutants ^37^. We conclude that, although Pak2 modulates polarized secretion, this is not its main function during fusion.

Pak1 acts redundantly to Pak2 during fusion by also negatively regulating the CIP, as shown by the increased phospho-MAPK^Pmk1^ in *pak1-ts* cells. Nevertheless, Pak1 inactivation does not raise Rho1-GTP levels at the fusion site, suggesting a distinct mode of action. However, several observations during mitotic growth have linked Pak1 to Rho signaling, as it delays activation of Rho1 during cytokinesis ^52^, phosphorylates the GAP^Rga8^ ^35^ but does not strongly modulate the CIP MAPK pathway ^16^. Since MAPK^Pmk1^ deletion does not restore fusion to PAK mutants, modulation of its activity does not appear critical for cell fusion regulation. We interpret its increased phosphorylation in *pak1-ts* as reflecting the activation of an upstream CIP component, perhaps another Rho GTPase.

### Cell wall biosynthesis driving fusion pore closure

A particularly intriguing feature of PAK mutants is their transient fusion phenotype ^1^, where a fusion pore is created in both cell wall and plasma membrane, allowing cytosol continuity, but is subsequently re-sealed. We understand this phenotype as an intermediate outcome between complete fusion failure and slow fusion pore expansion, due to overactive cell wall biosynthesis. Indeed, cryoSXT demonstrates that the cell wall has been rebuilt. Interestingly, *S. cerevisiae fig2Δ* cells, in which PKC is not suppressed at the fusion site, also exhibit transient fusion events ^63^, suggesting that unimpeded wall repair drives pore re-sealing across organisms. Thus, cell wall synthesis may exert sufficient force to promote plasma membrane fission. This echoes studies of cytokinesis, which have shown that wall synthesis can drive division site closure in absence of the actomyosin contractile ring ^68,69^. Thus, directionality of cell fusion is not guaranteed by membrane fusion but ensured by the PAK-dependent active repression of cell wall forces that would otherwise reverse it.

## Materials and Methods

### Strain construction

Standard genetic manipulation techniques were used for fission yeast *S. pombe* strains construction, by transformation or tetrad dissection. All strains, plasmids and primers are available on Tables S1, S2 and S3 respectively. The tagging of Wsc1, Pck1, Pck2 and Pmk1 by sfGFP, and the deletion of Rho4 were performed at the endogenous native locus by PCR-based homologous recombination ^70^ using the pFA6a plasmids pSM1538 or pSM644 as template and the primers listed on table S3, which amplified a cassette containing two 78nt-long homology regions at C-terminus of the ORF (or at 5’UTR for *rho4Δ*) and in the 3’ UTR respectively, the sfGFP ORF and/or a KanMx resistance cassette. The Myo52-mScarlet-I was obtained in a similar way and was previously described in ^4^. The Pak2 truncations and point mutations were performed by modifying the pFA6a plasmid pSM4231. After linearization, those constructs replaced the native Pak2 locus. Mutation of the PH domain basic residues K33, K45 and K58 to alanine is predicted to prevent phosphoinositide binding based on structural analysis of other PH domains ^71^. The mutation of H137 and H140 to leucine (HHLL), which is predicted to block Cdc42 binding, and the K343A point mutation, predicted to abolish kinase activity, were described previously ^22^. Plasmids pSM2951, pSM2952 and pSM3191 were constructed using the Single Integration Vector system described in ^72^. The split-GFP system comprises a SynZip4-GFP1-10 fragment (pSM2952) under the *tdh1* promoter at the *his5* locus and a SynZip3-GFP11 construct (pSM2951) under the *act1* promoter at the *ade6* locus. The RBD2-sfGFP construct (pSM3191) comprises the Pck2 promoter and its N-terminal RBD domain (aa 1-381) fused to sfGFP and targeted for genomic integration at the *ade6* locus.

### Media and growth conditions

For live microscopy, the method described in ^73^ was followed, with modifications. Cells were initially grown overnight in Minimal Sporulation Liquid (MSL) media with glutamate as the main nitrogen source (MSL+E) at 30°C until an OD_600_ between 0.8 and 1.0. Then cells were washed 3 times with MSL without nitrogen (MSL-N), adjusted to an OD_600_ of 1.5 and grown for 2-3 more hours at 30°C. In the case of split-GFP system, heterothallic *h+* and *h-* cells were mixed 1:1 right before MSL-N washes. Cells were then deposited on top of 2% electrophoresis-grade agarose pads of MSL-N with or without the addition of 1M Sorbitol covered with a coverslip sealed with VALAP (1:1:1 vaseline/lanolin/paraffin). Pads were either incubated for 24h at 25°C, 30°C or 32°C, to quantify the mating outcome, or, after letting them rest for 30’, directly used to acquire overnight timelapse imaging at room temperature.

### Live microscopy

24h post-starvation snapshots and timelapse imaging (frames every 5’) were performed with a Nikon ECLIPSE Ti2 inverted microscope operated by the NIS Elements software (Nikon). Images were acquired using a Plan Apochromat Lambda 100X 1.45 NA oil objective and recorded using Prime BSI express sCMOS camera from Teledyne Vision Solutions. Autofocusing was performed using the Nikon Perfect Focus System for long term movies with multipoint acquisition on single z-plane (xy pixel size 0.65 x 0.65 μm). Exposure time and laser intensity were kept constant for every set of tagged protein imaging and each set subjected to 3-color imaging using fluorescence excitation at 405/488/561nm with the Lumencor SpectraX light engine (Chroma) in addition to differential interference contrast (DIC) imaging with transmitted light. Images from Fig 1C, 2A, 2D-G that were obtained with a DeltaVision platform (Applied Precision) composed of a customized inverted microscope (IX-71; Olympus), a UPlan-Apochromat 100× /1.4 NA oil objective, a camera (CoolSNAP HQ2; Photometrics or 4.2Mpx Prime BSI sCMOS camera; Photometrics), and a color combined unit illuminator (Insight SSI 7; Social Science Insights). Images were acquired using softWoRx v4.1.2 software (Applied Precision). The Alexa Quad (DAPI/FITC/A594/Cy5) dichroic mirror filter set was used to record BFP (ex 390/18, em 435/48), RFP (ex 575/25 em 632/60), and GFP (ex 475/20 em 525/50) signals, with additional reference image taken in differential interference contrast (DIC) for each time point. All image visualization and processing were done with ImageJ/Fiji software (NIH, Bethesda, USA). For figure 3C, due to the relatively low signal of the glucanases, imaging was performed on live cells using single-frame acquisitions instead of timelapse. After N starvation for 4h, cells were mounted on MSL-N pads as described above. Imaging was carried out using a Zeiss LSM980 confocal microscope equipped with an Airyscan2 detector on an inverted Axio Observer 7 platform with 63x/1.40 NA oil immersion objective. Excitation was performed with 488 nm and 561 nm laser lines. Images were acquired at a resolution of 1863×1863 pixels (pixel size ∼0.04 μm), with 1.7x digital zoom, 16-bit depth, line averaging of 2, and a scan speed of 5. Unless otherwise stated, scale bars represent 5µm.

### Mating quantification

Mating outcomes were quantified on the 24h post-starvation snapshots. Only the cells engaged in mating were considered. For the Complete Fusion (CF) category, every fused pair and zygote containing four spores were counted as two CF cells. Single cells, paired and with a shmoo shape, without spores were counted as No Fusion (NF). Single cells, paired and with a shmoo shape, containing spores, a hallmark of haploid meiosis, were quantified as Transient Fusion (TF). These cells could either be paired to another non-fused cell or to a completely fused pair, in a 3-partner mating event, as previously described ^1^. For mating-associated cell lysis (LY), only paired (shmoo-shape) lysed cells were quantified. The pairing efficiency in Fig 6E, displays the proportion of the cell population engaged in mating, irrespective of their mating outcome. In all cases, three independent replicates were analyzed, with n ≥ 400 cells per replicate (n ≥ 600 for pairing efficiency), and statistics were performed (paired-t-test or Wilcoxon paired test) on the complete fusion fraction.

### Fluorescence-loss in photobleaching (FLIP)

This was performed on a spinning disk microscope composed of an inverted microscope (DMI4000B; Leica) equipped with an HCX Plan Apochromat 100×/1.46 NA oil objective, a CSU22 real-time confocal scanning head (Yokogawa Electric Corporation), solid-state laser lines (Visitron LaserModule 1870), and an electron multiplying charge-coupled device camera (iXon Life 897 Back-Illuminated EMCCD Camera). Photobleaching and timelapse image acquisition were controlled by the VisiView Premier GOLD Image acquisition Software (Visitron). Heterothallic cells expressing the split-GFP system and mixed 1:1 were N starved and plated on a MSL-N pad for 24h at 25°C. An initial image was acquired in both GFP and brightfield to identify GFP-positive pairs and distinguish complete fusion (CF) from transient fusion (TF) based on visualization of the cell wall in the brightfield image. After photobleaching laser calibration, 2 µm-diameter circle area were drawn on a cytosolic region of one of the cells of the selected GFP-positive pair. The region was bleached over 29 cycles every 3’’, with 1 or 3 frame acquired before first bleaching in the *pak2*Δ vs the *pak2Δ pak1-ts* respectively. For quantification, 5×5 px areas were drawn i) in the bleached region, ii) in the unbleached partner cell, iii) in an unbleached control cell to measure photobleaching. With an in-house MATLAB macro, the photobleaching decay was calculated, then photobleaching-corrected profiles were normalized to the value before bleaching. The mean of the bleached spot and the CF examples were plotted together with each TF individual profile, to classify them as closed, if no decay was detected, or open if the decay was similar to the one of the CF examples.

### Pore expansion quantification

Timelapses of *h90* strains expressing Wsc1-sfGFP were taken (images every 5’). Time was relative to fusion time, with t = 0 defined as the first frame after the entry of the mTagBFP2 expressed from the P cell into the M cell. Background subtraction was applied to the images, followed by alignment to compensate for stage drift using an in-house ImageJ macro ^4^ (https://github.com/Valentine-Thomas/FusionFocus-Mapping/blob/master/Image_Processing/Centroid-detection-3channels-method3.ijm) On the selected pairs, a 20px-wide line was drawn across the fusion pore, labelled by the position of the fusion focus marker Myo52-mScarlet-I at t = -10’. The red channel intensity at t = -10’ and the green channel intensity at t = +5’, +15’, +25’ and +35’ were obtained. Using an in-house R macro, the profiles were aligned and plotted setting x = 0 as the max intensity of Myo52 at t = - 10’. Then those profiles were used to build double-gaussian models (in house-MATLAB macro), and the peak distance was defined as the distance of the max of each of the two gaussians. A linear regression was used (using an R macro) to calculate the pore expansion rate.

### Fusion time measurement

Fusion time was calculated, from 5’ frame rate timelapses, as the time gap between the first frame in which a clear Myo52-labelled fusion focus is identified and the first frame after the P cell cytosolic marker transfers inside the M cell.

### Fluorescence intensity over time

To plot the intensity over time in Fig 2B and 5B, the fusing pairs were identified on squared ROI comprising the full pair. Images were first treated as in ^4^ to subtract background, perform photobleaching correction, crop out every ROI, align in space to compensate for stage drift and in time by selecting the -1h to +2h window. Then a segmented, 20px-wide line was drawn along the cell-cell axis in each aligned cropped movies and a kymograph was obtained. On the kymograph, a 5px-wide straight line was drawn along the time axis at the location of the fusion focus. The intensities from the red and green channels were obtained, normalized to min and max in Fig 2B or to the cytosol in Fig 5B and plotted using R studio.

### Fluorescence parameters at fusion focus

To calculate the focus to cytosol ratio in Fig 2E, the timelapse images were first background-subtracted and photobleaching-corrected. On each of the selected pairs, at the t = -5’ frame, a 5×5 px area was drawn on the Myo52-mScarlet-I labelled fusion focus, and two freehand-shaped ROIs were drawn on the cytosolic areas of the two partner cells. Then the intensities on the green channel were obtained, and the intensity at the fusion focus was divided by the mean of the intensities of the two cytosolic areas. To calculate the full width at half-maximum (FWHM) (Fig 2F, 3B) and area under the curve (Fig 3B, S1B), timelapses were acquired on a Nikon ECLIPSE Ti2 inverted microscope, and image stacks were corrected for photobleaching and stage drift as described for the pore expansion calculations above. On the identified cell pairs, a 20-pixel-wide straight line was drawn across the fusion focus 10’ before fusion (or 5’ in Fig 2F) and fluorescence intensity profiles were extracted and fitted in MATLAB to a Gaussian function *f*(*x*) = *A*exp[−(*x* − *μ*)^2^/(2*σ*^2^)], where *A* is the amplitude, *μ* the center, and *σ* the standard deviation. FWHM values were derived from the fitted *σ* using FWHM = 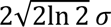, and the area under curve was obtained as the integrated intensity by applying MATLAB’s *trapz* function to each profile to approximate the integral as the sum of trapezoidal segments between successive data points.

### Fluorescence parameters at contact site

For the measurements in Fig 5C-I timelapse images were first aligned to compensate for stage drift. On the selected pairs, a 20px-wide line was drawn across the fusion pore, labelled by the position of the fusion focus marker Myo52-mScarlet-I at t = -10’. The red channel intensity at t = -10’ and the green channel intensity at t = +25’ were obtained, and profiles were aligned to the max of Myo52 as described at the pore expansion method above. Additionally, a freehand-shaped ROIs were drawn on the cytosolic area of one of the partners at -10’, and the corresponding mean values for the green channel were obtained. On the aligned profiles, a second in-house R macro subtracted the background of both the profiles and the cytosol values, and calculated the corresponding signal to cytosol ratio for every value on the profile, generating the plots over the contact site in Fig 5D,E. The values on Fig 5 F-I were extracted from these plots, defined as the mean of the signal around the central 1µm (from -0.5 to 0.5 µm).

### Protein extraction for western blotting

After two sequential dilutions, cells were grown overnight in MSL+E at 30°C until an OD_600_ between 0.8 and 1.0 and a final volume of 150mL (or 70mL in Fig S1C). After separating 30mL as the vegetative sample, fixing them with a final 10% trichloroacetic acid (TCA) and keeping them on ice, the remaining cells were washed by vacuum filtration using sterile cellulose nitrate filters (0.45µm, Sartorius 11306-47-N) and eluted with 20mL (or 5mL in S1C) of MSL-N. Cultures were adjusted to a final OD_600_ of 1.5 in final volume of 50 mL (or 20mL in S1C) in MSL-N. The heterothallic *h+* and *h-* cells were mixed 1:1 post starvation, except in Fig S2H, where all 3 strains were kept on different flasks, and only mixed upon TCA fixation. Cells were immediately cultured at 25°C, being this moment the time 0 post-starvation. At the indicated timepoints (and at 5h post-starvation in Fig S1C), 18mL of culture mix (or 9 + 9mL in Fig S2H) were taken and mixed with 2mL of TCA for a final 10% concentration and kept on ice until all samples were fixed. Protein extracts were obtained following the protocol described in ^74^. The lysis buffer was complemented with complete protease inhibitor cocktail tablets (Roche, 11836145001)) and PhosSTOP tablets (1 in 10 ml, Roche, 04 906 837 001) to inhibit phosphatase activity and preserve MAPK phosphorylation. After washing, cells were resuspended in 300 μL lysis buffer. Cell were lysed using a MagNA Lyser bead beater (Roche) for four cycles of 45 s at 6,500 rpm (with 1’ beak between cycles). The lysate supernatant was collected, and protein concentration was measured by Bradford assay. Samples were further denatured at 65 °C in 4× NuPAGE LDS sample buffer (Invitrogen, NP0007) for 15’, followed by β-Mercaptoethanol (1 μL per 20 μL sample) addition and incubated 10’ at RT before SDS-PAGE.

### Western blotting

Adjusted amounts of protein were loaded onto 4%–12% acrylamide gels (GenScript) and electrophoresed at 120 V for 1h 30’ using commercial Tris-MOPS-SDS running buffer (GenScript). Proteins were transferred onto a nitrocellulose membrane (Cytiva) using a homemade transfer buffer (50 mM Tris base, 38 mM glycine, 1% SDS, 20%ethanol) at 110 V for 1h 30’. PageRuler (Thermo, 26619) was used as protein ladder. Membranes were incubated overnight with anti-phospho-p44/42 MAPK (rabbit, Cell Signaling Technology, 4370, at 1:1000), anti-GFP (mouse, roche 11814460001 at 1:3000), or anti-Actin (mouse, abcam, ab8224, 1:5000) as primary antibodies. followed by the secondary antibodies (1:10000 dilution) anti-rabbit IRDye 800CW or anti-mouse IRDye 800CW and IRDye 680RD (LICORbio). The membrane was developed using the Odyssey imaging system (LI-COR), and the results were quantified using imageJ as described in ^37^. All uncropped raw image files are available in Data S1-5.

### Cryo-Soft-X-ray Tomography (cryo-SXT)

A mating mixture of P cells (*h+*) expressing SynZip4–GFP(1–10), and M cells (*h−*) expressing SynZip3–GFP(11) and Myo52-mScarlet-I was nitrogen starved. 30 OD_600_ of cells were resuspended in 1mL MSL-N and 50µL drops were plated on MSL-N plates at 25°C for 5h (WT) or 12h (*pak2Δ pak1-ts*). After a fluorescence microscopy check, cells from two of the drops were harvested, resuspended in 4mL MSL-N liquid media and adjusted to an OD_600_ = 0.9. Cells were cryo-fixed by plunge-freezing using Leica EM GP2–Automatic Plunge Freezer, by placing 3.5µL cell suspension on Quantifoil R2/1 200 mesh Copper Grids, with the addition of 1µL 100nm gold nanoparticles suspension (3.60×10^8^ particles/mL, EMGC100, BBI Group, Cardiff, UK), with 3s back blotting, on a chamber temperature of 23°C and 99% humidity. The fixed grids were clipped and then imaged by cryo-fluorescence microscopy using a Leica cryoThunder microscope (50X), and shipped under cryogenic conditions to ALBA synchrotron. The cryo-fluorescence grid images were deeply annotated to identify all the post-fused (split-GFP positive) pairs before SXT image acquisition. Clipped grids were transferred under cryogenic conditions to the MISTRAL beamline (ALBA light source) at ALBA Synchrotron ^75,76^. Tomographic data were collected at 520 eV, irradiating the samples for 1-2’’ per projection. Projection images obtained at different sample orientations were computationally combined to produce a three dimensional (3D) reconstruction, permitting the 3D representation of the subcellular ultrastructure of whole cells ^77^. A tilt series was acquired for each cell area using an angular step of 1 ◦ on a ±70 ◦ range with a Fresnel Zone plate (FZP) of 25 nm outermost zone width and an effective pixel size of 11 nm. Each transmission projection image of the tilt series was normalized using flat-field, accounting for the exposure times, as well as the machine current. Wiener deconvolution considering the experimental impulse response of the optical system was applied to the normalized data in order to increase the image quality ^78^. Finally, the Napierian logarithm was used to reconstruct the linear absorption coefficient (LAC). The resulting stacks were then loaded into IMOD software ^79^ and the individual projections were aligned to the common tilt-axis using the gold particles of 100nm as markers. Subsequently, the aligned stacks were reconstructed with algebraic reconstruction techniques (ART) ^80^. The visualization and segmentation of the volumes were carried out using Amira 3D software (ThermoFisher) and volume scenes and movies were generated using Chimera X ^81^.

The tomograms and their corresponding fluorescence information were used to classify them as either post-fusion open pore or post-fusion resealed. Pore width is defined as the maximum distance between the two inner surfaces of the cell wall across the volume of the fusion pore.

### Statistics

Stacked bar and violin plots were prepared on GraphPad Prism, shadowed curve profiles were built on R studio. After a normality test check, two-tailed t-test, (or Mann-Whitney non-parametric test, if not normal) were run on GraphPad Prism. For paired sample comparison, a paired two-tailed t-test or a non-parametric paired Wilcoxon test were performed.

## Author contribution

The project was designed by JMCS and SGM. JMCS performed all experiments and analyses, except Fig 3 B-C, which were performed by SS. AJPB helped cryo-SXT data acquisition and segmentation. SGM and JMCS acquired funding and wrote the paper.

## Acknowledgements

We thank Aleksandar Vještica and Léo Franchi (UNIL, Lausanne) and Juan Carlos Ribas and Pilar Pérez (IBFG, Salamanca) for strains, Sushila Gordon-Lennox and Laetitia Michon for technical help, Olivia Muriel-López for preliminary exploration of the resealing pore ultrastructure, Valentine Thomas, Wanlan Li and Ingrid Billault-Chaumartin for help with data analysis code, the Dubochet Center for Imaging at UNIGE for help in cryoSXT sample preparation and members of the lab for discussion. This work was supported by an EMBO postdoctoral fellowship (ALTF 351-2021) to JMCS, and a Swiss National Science Foundation grant (310030_207909) and ERC Advanced Grant (SexYeast; 101019630) to SGM. Cryo-SXT was performed at the MISTRAL beamline at ALBA Synchrotron funded by the 2024-II call (2024028120).

---

**Figure S1.**
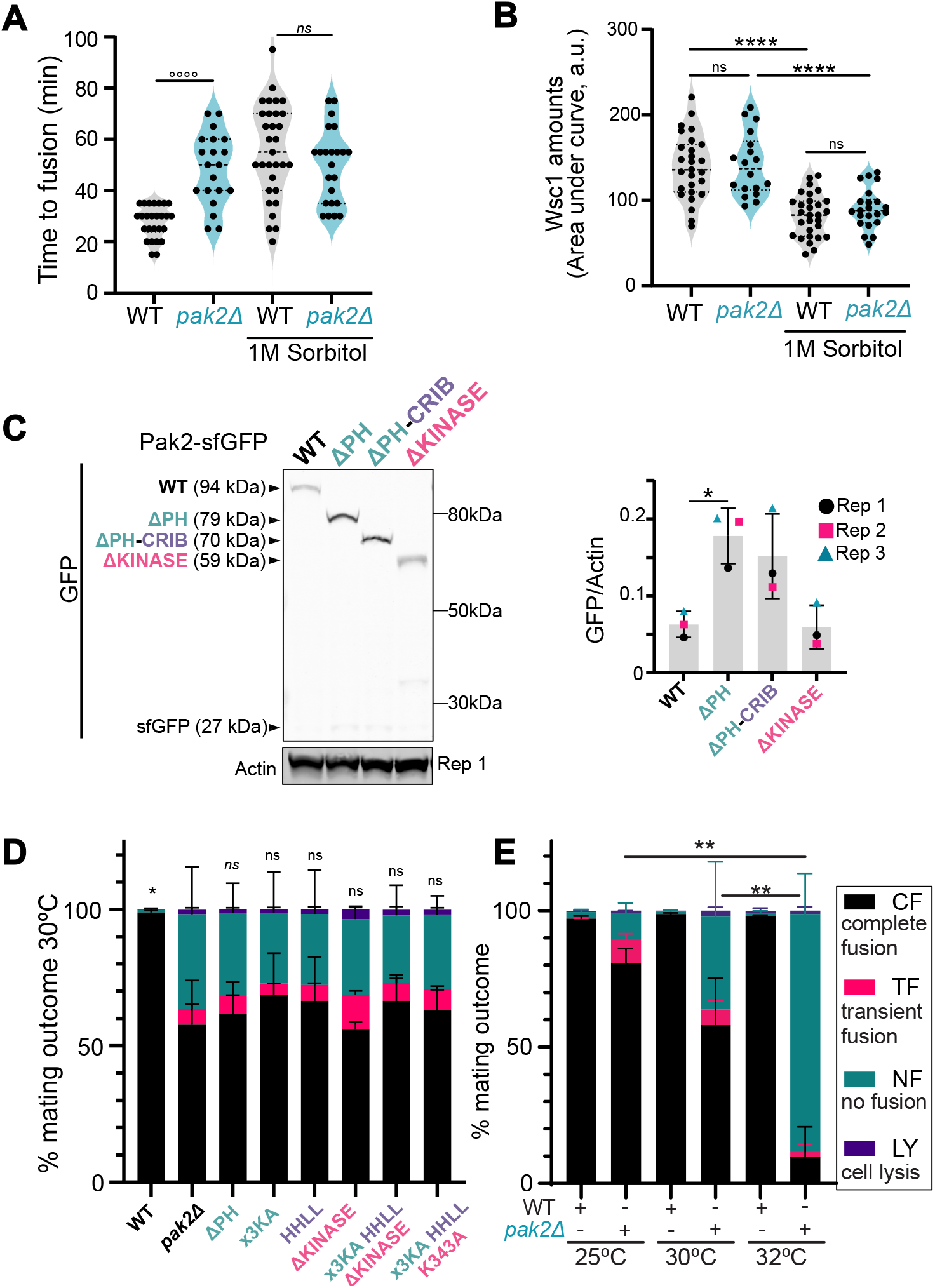
Fusion time measurements, Wsc1 intensity levels, stability and phenotype of *pak2* mutants, effect of temperature on *pak2Δ*. (Related to Fig 1, 2 and 4) (**A**) Measurement of fusion time in WT vs *pak2Δ* in absence (n = 26, 20) or presence of 1M sorbitol (n = 31, 23). Statistics show Mann-Whitney tests, with °°°° = p<0.0001, *ns* = p≥0.05. (**B**) Amounts (area under curve) of Wsc1 at fusion site at -10’ in WT vs *pak2Δ* in absence (n = 27, 18) or presence of 1M sorbitol (n = 29, 22). Statistics show t-tests, with **** = p<0.0001, ns = p≥0.05. (**C**) Western blot and quantification of indicated Pak2-sfGFP construct in *h90* strains. Statistics show t-test, with * p<0.05. (**D**) Frequency of mating outcomes of indicated strains. Statistics show paired t-test on the complete fusion frequency vs *pak2Δ*, with * = p<0.05, **= p<0.005, ns = p≥0.05 .Italized *ns* p≥0.05 indicate a non-parametric paired Wilcoxon test. (**E**) Frequency of mating outcomes of indicated strains and temperatures. Statistics show paired t-test on CF frequency with **= p<0.005.

**Figure S2.**
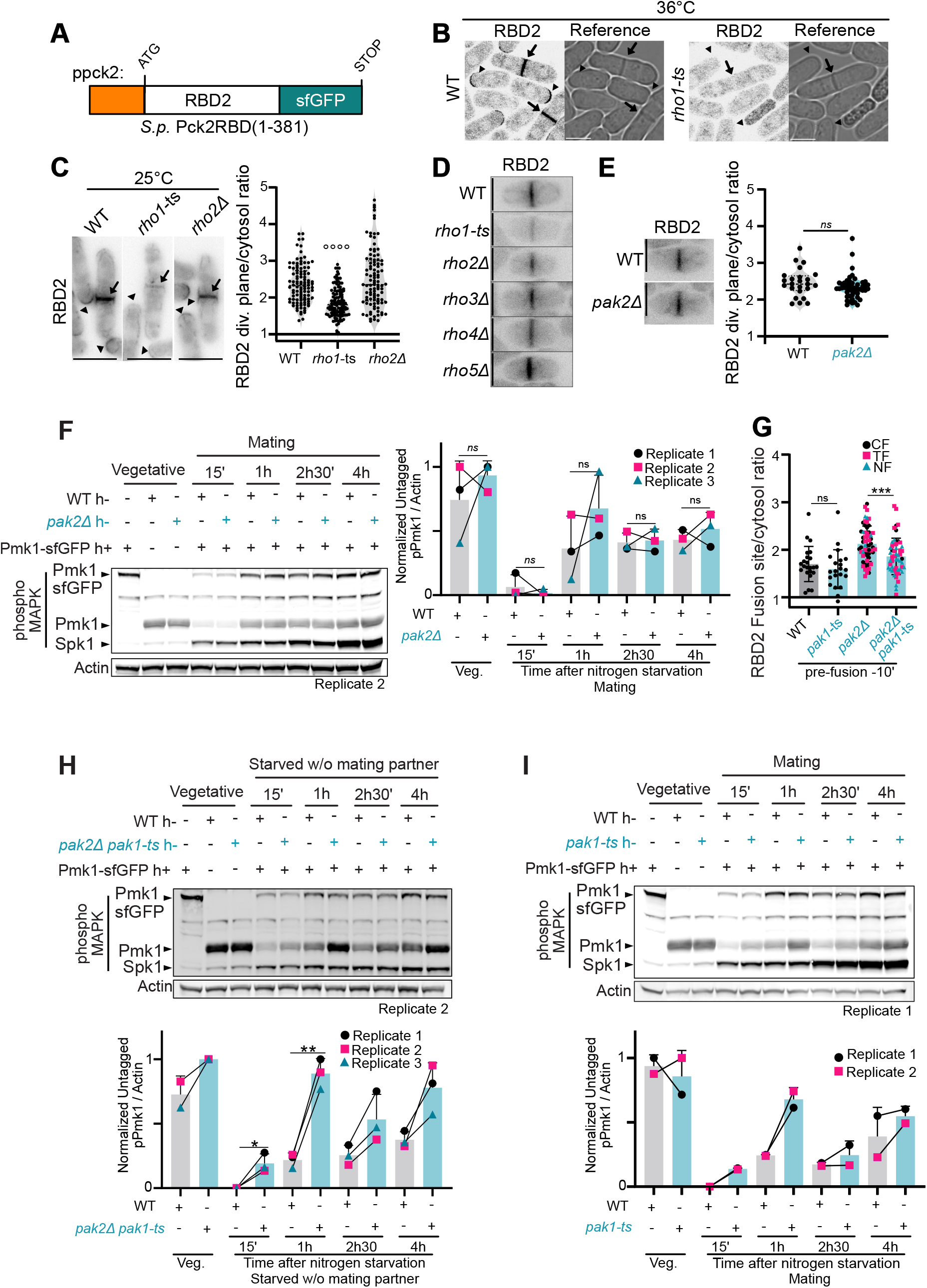
Characterization of RBD2-sfGFP, and effect of pak2 on the phosphorylation levels of Pmk1. (Related to Fig 5) **(A)** Scheme of the RBD2-sfGFP probe. (**B-E**) Localization of RBD-sfGFP in indicated strains grown in rich media after 1h incubation at 36°C (B) at 25°C (C) or upon nitrogen starvation at 25°C (D-E). Arrows = division plane, arrow heads = cell tips. The plots show quantification of the division plane to cytosol ratio of RBD2 (n =104, 124, 100 in (C); n = 24, 49 in (E)). Statistics show Mann-Whitney tests, with °°°° = p<0.0001 or *ns* = p≥0.05, vs the WT control. (**F**) Western blot of phospho-Pmk1 levels as in Fig 5J for WT and *pak2Δ*. (**G**) RBD2 fusion site to cytosol ratio pre-fusion in WT, *pak1-ts*, *pak2Δ* and *pak2Δ pak1-ts* pairs, measure as in Fig 5F. Statistics show t-test with *** p<0.001 and ns p≥ 0.05. (**H**) Western blot of phospho-Pmk1 levels during starvation as in Fig 5J, except that strains were starved separately and only mixed after TCA fixation. (**I**). Western blot of phospho-Pmk1 levels as in Fig 5J for WT and *pak1-ts*. Error bar represent standard deviation. In (F, H, I), statistics show paired t-test with * p<0.05, ** p<0.005, ns = p≥0.05 Italized *ns* p≥0.05 indicate a non-parametric paired Wilcoxon test.

**Figure S3.**
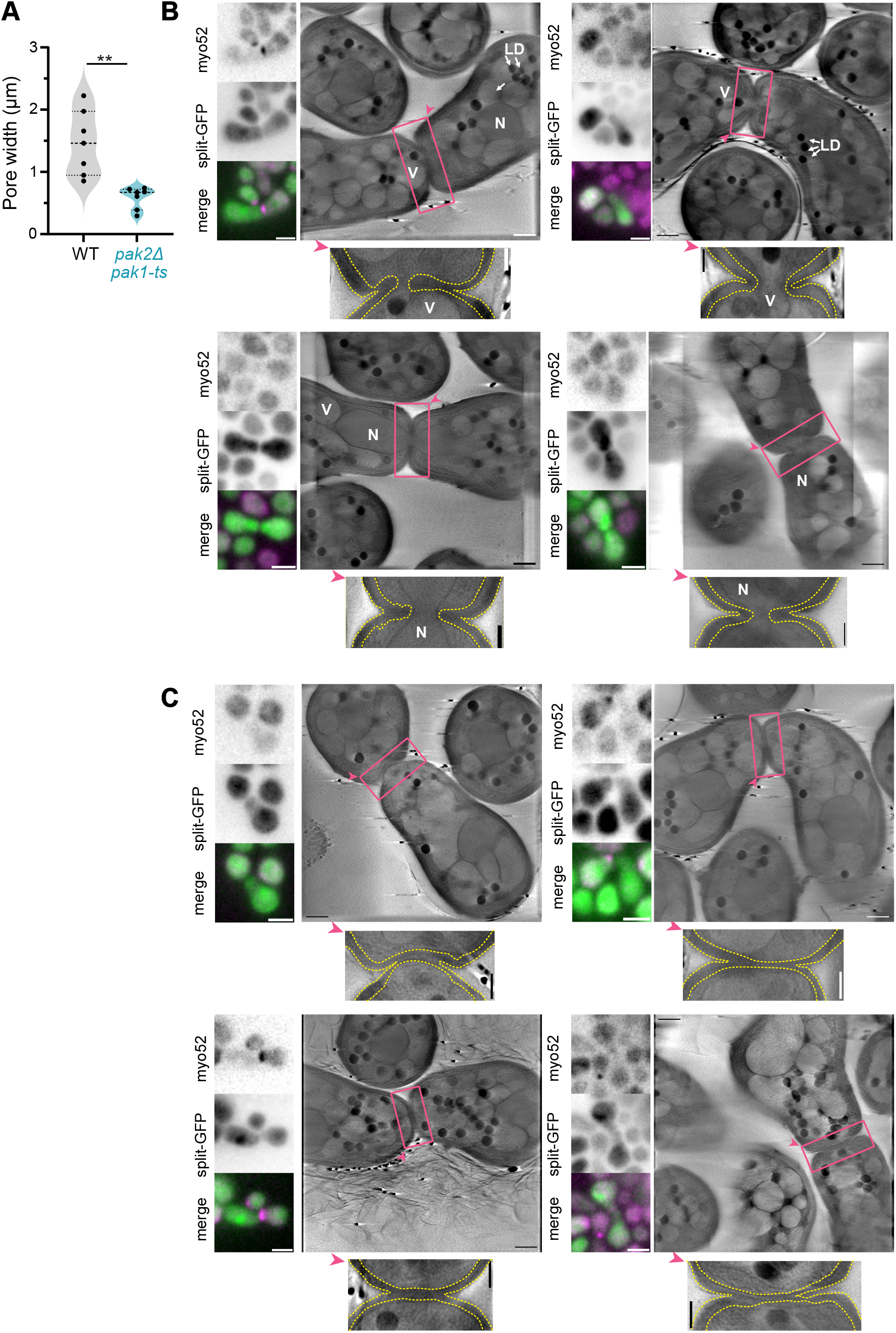
Other cryo-SXT images of post-fusion PAK mutants. (Related to Fig 7) (**A**) Pore width measured from cryo-SXT tomograms of WT and *pak2Δ pak1-ts* completely fused pairs (n = 7, 7). Statistics show t-test with ** p>0.005. (**B-C**) Correlative light-cryo-SXT on a double mutant *pak2Δ pak1-ts* strain pairs with four examples each of narrow opened pore, plugged by a vacuole or nucleus (**B**) or transient fused and resealed pairs (**C**). On the left of each subpanel, the corresponding fluorescence microscopy of the selected pair. The pink arrowhead indicates orientation of the enlarged region. Scale bars of fluorescence panels = 5 µm, full cryo-SXT = 1 µm, and enlarged region = 0.5 µm. N = nucleus, V= vacuole, LD = lipid droplets. The cell wall is indicated as a yellow dotted line (enlarged regions).

**Movie S1. Full-volume cryo-SXT tomogram of WT fused cell pair. (Related to Fig 7**) Full volume of the cryo-SXT tomograms displayed in Fig 4B showing a WT fused pair. Nucleus = pink volume, Cell wall = yellow surface.

**Movie S2. Full-volume cryo-SXT tomogram of *pak2Δ pak1-ts* fused cell pair. (Related to Fig 7)**

Full volume of the cryo-SXT tomograms displayed in Fig 4C showing a *pak2Δ pak1-ts* pair with a narrow pore plugged with a vacuole. Nucleus = pink volume, Vacuole = violet volume, Cell wall = yellow surface.

**Movie S3. Full-volume cryo-SXT tomogram of *pak2Δ pak1-ts* transiently fused cell pair, with rebuilt cell wall. (Related to Fig 7)**

Full volume of the cryo-SXT tomograms displayed in Fig 4d showing a *pak2Δ pak1-ts* pair with a re-sealed pore and apparently intact cell wall. Nuclei = pink volume, Cell wall = yellow and blue surfaces.

