## Supplementary material for "PAK-family kinases promote cell fusion irreversibility by preventing cell wall repair": Table S1

**Table S1: Strains used in this study**

Strains are indicated in the first Figure where they appear. The methods column indicates how they were obtained. Please refer to Tables S2 and S3 for details of the plasmids and primers.

| | Strain | Genotype | Method |
|---|---|---|---|
| | YSM4457 | h90 myo52-mScarlet-I:natMX lys3+:pmap3:mTagBFP2:termScAdh1:bleMX ade6+:ppck2:RBDpck2-sfGFP:bsdMX ura4+ leu1+ his+ | Cross JCS0389 x YSM4259 |
| | YSM4458 | h90 pak2Δ::hphMX myo52-mScarlet-I:natMX lys3+:pmap3:mTagBFP2:termScAdh1:bleMX ade6+:ppck2:RBDpck2-sfGFP:bsdMX ura4+ leu1+ his+ | Cross JCS0389 x YSM4259 |
| | YSM4459 | h90 orb2-34 myo52-mScarlet-I:natMX lys3+:pmap3:mTagBFP2:termScAdh1:bleMX ade6+:ppck2:RBDpck2-sfGFP:bsdMX ura4+ leu1+ his+ | Cross JCS0540 xYSM4457 |
| | YSM4460 | h90 orb2-34 pak2Δ::hphMX ade6+:ppck2:RBDpck2-sfGFP:bsdMX myo52-mScarlet-I:natMX lys3+:pmap3:mTagBFP2:termScAdh1:bleMX ura4+ leu1+ his+ | Cross JCS0120 x YSM4458 |
| | YSM4461 | h90 pak2-K343A-sfGFP:kanMX myo52-mScarlet-I:natMX lys3+:pmap3:mTagBFP2:termScAdh1:bleMX ura4+ leu1+ his+ | JCS0243 transformed with AfeI linearized pSM3028 |
| Figure 1 | YSM4462 | h90 orb2-34 pak2-K343A-sfGFP:kanMX myo52-mScarlet-I:natMX lys3+:pmap3:mTagBFP2:termScAdh1:bleMX ura4+ leu1+ his+ | Cross of YSM4461 x JCS0540 |
| | YSM4463 | h+ pak2Δ::hphMX his5+:ptdh1:SynZip4-GFP1-10:natMX ura4+ leu1+ ade6+ | cross YSM4479 x YSM4264 |
| | YSM4464 | h- pak2Δ::hphMX ade6+:pact1:SynZip3-GFP11:bsdMX ura4+ leu1+ his+ | cross JCS0057 x JCS0060 |
| | YSM4465 | h+ orb2-34 pak2Δ::hphMX his5+:ptdh1:SynZip4-GFP1-10:natMX ura4+ leu1+ ade6+ | Cross JCS0129 xJCS0040 |
| | YSM4466 | h- orb2-34 pak2Δ::hphMX ade6+:pact1:SynZip3-GFP11:bsdMX ura4+ leu1+ his+ | Cross YSM4464xJCS120 |
| | YSM4467 | h90 wsc1-sfGFP:kanMX myo52-mScarlet-I:natMX lys3+:pmap3:mTagBFP2:termScAdh1:bleMX ura4+ leu1+ ade6+ his+ | Cross JCS0799 x JCS0800 |
| | YSM4468 | h90 pak2Δ::hphMX wsc1-sfGFP:kanMX myo52-mScarlet-I:natMX lys3+:pmap3:mTagBFP2:termScAdh1:bleMX ura4+ leu1+ ade6+ his+ | Cross JCS0736 x YVT186 |
| | YSM4469 | h90 pak2-sfGFP:kanMX myo52-mScarlet-I:natMX lys3+:pmap3:mTagBFP2:termScAdh1:bleMX ura4+ leu1+ ade6+ his+ | JCS0005 was transformed with SpeI linearized pAV0761 |
| | YSM4470 | h90 pak2(129-589)-sfGFP:kanMX myo52-mScarlet-I:natMX lys3+:pmap3:mTagBFP2:termScAdh1:bleMX ura4+ leu1+ ade6+ his+ | YVV776 was transformed with AfeI linearized pSM3042 |
| | YSM4471 | h90 pak2-K33A,K45A,K58A-sfGFP:kanMX myo52-mScarlet-I:natMX lys3+:pmap3:mTagBFP2:termScAdh1:bleMX ura4+ leu1+ ade6+ his+ | JCS0243 transformed with AfeI linearized pSM3067 |
| | YSM4472 | h90 pak2(206-589)-sfGFP:kanMX myo52-mScarlet-I:natMX lys3+:pmap3:mTagBFP2:termScAdh1:bleMX ura4+ leu1+ ade6+ his+ | YVV776 was transformed with AfeI linearized pSM3043 |
| | YSM4473 | h90 pak2-H137L,H140L-sfGFP:kanMX myo52-mScarlet-I:natMX lys3+:pmap3:mTagBFP2:termScAdh1:bleMX ura4+ leu1+ ade6+ his+ | YVV776 was transformed with pSM3057 |
| Figure 2 | YSM4474 | h90 pak2(1-279)-sfGFP:kanMX myo52-mScarlet-I:natMX lys3+:pmap3:mTagBFP2:termScAdh1:bleMX ura4+ leu1+ ade6+ his+ | YVV776 was transformed with AfeI linearized pSM3044 |
| | YSM4520 | h90 pak2(206-589)-K343A-sfGFP:kanMX myo52-mScarlet-I:natMX lys3+:pmap3:mTagBFP2:termScAdh1:bleMX ura4+ leu1+ ade6+ his+ | YVV776 was transformed with AfeI linearized pSM3049 |
| | YSM4475 | h90 pak2(1-279)-K33A,K45A,K58A,H137L,H140L-sfGFP:kanMX myo52-mScarlet-I:natMX lys3+:pmap3:mTagBFP2:termScAdh1:bleMX ura4+ leu1+ ade6+ his+ | JCS0243 was transformed with AfeI linearized pSM3166 |
| | YSM4476 | h90 pak2-K33A,K45A,K58A,H137L,H140L,K343A-sfGFP:kanMX myo52-mScarlet-I:natMX lys3+:pmap3:mTagBFP2:termScAdh1:bleMX ura4+ leu1+ ade6+ his+ | JCS0243 transformed with AfeI linearized pSM3097 |
| | YSM4477 | h90 pak2(206-589):kanMX myo52-mScarlet-I:natMX lys3+:pmap3:mTagBFP2:termScAdh1:bleMX ade6+:ppck2:RBDpck2-sfGFP:bsdMX ura4+ leu1+ his+ | YSM4457 was transformed with AfeI linearized pSM3006 |
| | YSM4478 | h90 pak2(206-589)-K343A:kanMX myo52-mScarlet-I:natMX lys3+:pmap3:mTagBFP2:termScAdh1:bleMX ade6+:ppck2:RBDpck2-sfGFP:bsdMX ura4+ leu1+ his+ | YSM4457 was transformed with AfeI linearized pSM3023 |
| | YSM4275 | h90 myo52-GFP:kanMX lys3+:pmap3:mTagBFP2:termScAdh1:bleMX ura4+ leu1+ ade6+ his+ | (37) |
| | YSM4521 | h90 myo52-GFP:kanMX pak2Δ::hphMX lys3+:pmap3:mTagBFP2:termScAdh1:bsdMX ura4+ leu1+ ade6+ his+ | YSM3931 was transformed with SpeI linearized pSM2955 |
| | YSM4269 | h90 fus1-sfGFP:kanMX lys3+:pmap3:mTagBFP2:termScAdh1:bsdMX ura4+ leu1+ ade6+ his+ | (37) |
| | YSM4522 | h90 fus1-sfGFP:kanMX pak2Δ::hphMX lys3+:pmap3:mTagBFP2:termScAdh1:natMX ura4+ leu1+ ade6+ his+ | SAS113 was transformed with SpeI linearized pSM3871 |
| | YSM4267 | h90 ade6+:GFP-Ypt3:hphMX lys3+:pmap3:mTagBFP2:termScAdh1:bleMX ura4+ leu1+ his+ | (37) |
| | YSM4523 | h90 ade6+:GFP-Ypt3:hphMX pak2Δ::kanMX lys3+:pmap3:mTagBFP2:termScAdh1:natMX ura4+ leu1+ his+ | SAS150 was transformed with SpeI linearized pSM3871 |
| Figure 3 | YSM4304 | h90 exo70-GFP:kanMX lys3+:pmap3:mTagBFP2:termScAdh1:bsdMX ura4+ leu1+ ade6+ his+ | (37) |
| | YSM4524 | h90 exo70-GFP:kanMX pak2Δ::hphMX lys3+:pmap3:mTagBFP2:termScAdh1:bleMX ura4+ leu1+ his+ | Cross SAS105 x JCS1047 |
| | YSM2573 | h90 exg3-sfGFP:kanMX myo52-tdTomato:natMX ura4+ ade6+ leu1+ his+ | (7) |
| | YSM4525 | h90 exg3-sfGFP:kanMX pak2Δ::hphMX myo52-tdTomato:natMX ura4+ ade6+ leu1+ his+ | YSM2573 was transformed with AfeI linearized pAV0220 |
| | ysm2571 | h90 agn2-sfGFP:kanMX myo52-tdTomato:natMX ura4+ ade6+ leu1+ his+ | (7) |
| | YSM4526 | h90 agn2-sfGFP:kanMX myo52-tdTomato:natMX pak2Δ::hphMX ura4+ ade6+ leu1+ his+ | YSM2571 was transformed with AfeI linearized pAV0220 |
| | YSM4527 | h90 agn1-GFP::Kan myo52-tomato-natMX ura4+ ade6+ leu1+ his+ | Cross YSM1396 x YSM2547 |
| | YSM4528 | h90 agn1-GFP::Kan myo52-tomato-natMX pak2Δ::hphMX ura4+ ade6+ leu1+ his+ | YSM4527 was transformed with AfeI linearized pAV0220 |
| | YSM4529 | h90 pck1-sfGFP:kanMX myo52-mScarlet-I:natMX lys3+:pmap3:mTagBFP2:termScAdh1:bleMX ura4+ leu1+ ade6+ his+ | Cross JCS0243 x YLF144 |
| | YSM4530 | h90 pak2Δ::hphMX pck1-sfGFP:kanMX myo52-mScarlet-I:natMX lys3+:pmap3:mTagBFP2:termScAdh1:bleMX ura4+ leu1+ ade6+ his+ | Cross JCS0243 x YLF144 |
| | YSM4531 | h90 pck2-sfGFP:kanMX myo52-mScarlet-I:natMX lys3+:pmap3:mTagBFP2:termScAdh1:bleMX ura4+ leu1+ ade6+ his+ | YVV776 was transformed with a PCR cassette using pSM1538 as template and primers osm8031 and 8032 |
| | YSM4532 | h90 pak2Δ::hphMX pck2-sfGFP:kanMX myo52-mScarlet-I:natMX lys3+:pmap3:mTagBFP2:termScAdh1:bleMX ura4+ leu1+ ade6+ his+ | JCS0243 was transformed with a PCR cassette using pSM1538 as template and primers osm8031 and 8032 |
| Figure 5 | YSM4393 | h- ura4+ ade6+ leu1+ his+ | WT strain |
| | YSM4533 | h+ bgs4Δ::ura4+ GFP-bgs4:leu1+ myo52-mScarlet-I:natMX lys3+:pmap3:mTagBFP2:termScAdh1:bleMX ade6+ his+ | Cross JCS0849 x YMB357. bgs4Δ::ura4+ GFP-bgs4:leu1+ described in (55) |
| | YSM4534 | h- pak2Δ::hphMX ura4+ ade6+ leu1+ his+ | Cross JCS0057 x JCS0060 |
| | YSM4535 | h+ bgs4Δ::ura4+ GFP-bgs4:leu1+ pak2Δ::hphMX myo52-mScarlet-I:natMX lys3+:pmap3:mTagBFP2:termScAdh1:bleMX ade6+ his+ | Cross JCS0243 x JCS0849. bgs4Δ::ura4+ GFP-bgs4:leu1+ described in (55) |
| | YSM4536 | h- orb2-34 pak2Δ::hphMX ura4+ ade6+ leu1+ his+ | Cross YSM4464 x JCS0120 |
| | YSM4537 | h+ pmk1-sfGFP:kanMX myo52-mScarlet-I:natMX ura4+ leu1+ ade6+ his+ | YSM4218 was transformed with a PCR cassette using pSM1538 as template and primers osm8035 and 8036 |
| | YSM4538 | h90 rho1-596:natMX myo52-mScarlet-I:natMX lys3+:pmap3:mTagBFP2:termScAdh1:bleMX ade6+:ppck2:RBDpck2-sfGFP:bsdMX ura4+ leu1+ his+ | Cross JCS0441 x JCS0442, rho1-596:natMX comes from PPG6840 (a gift from Pilar Perez) |
| | YSM4539 | h90 rho1-596:natMX pak2Δ::hphMX myo52-mScarlet-I:natMX lys3+:pmap3:mTagBFP2:termScAdh1:bleMX ade6+:ppck2:RBDpck2-sfGFP:bsdMX ura4+ leu1+ his+ | Cross JCS0441 x JCS0442, rho1-596:natMX comes from PPG6840 (a gift from Pilar Perez) |
| | YSM4540 | h90 pck1Δ::kanMX myo52-mScarlet-I:natMX lys3+:pmap3:mTagBFP2:termScAdh1:bleMX ade6+:ppck2:RBDpck2-sfGFP:bsdMX ura4+ leu1+ his+ | osm8030 |
| | YSM4541 | h90 pck1Δ::kanMX pak2Δ::hphMX myo52-mScarlet-I:natMX lys3+:pmap3:mTagBFP2:termScAdh1:bleMX ade6+:ppck2:RBDpck2-sfGFP:bsdMX ura4+ leu1+ his+ | osm8030 |
| | YSM4542 | h90 orb11-59 ura4+ leu1+ ade6+ his+ | Cross JCS0237 x JCR544 (a gift from JC Ribas) |
| | YSM4543 | h90 orb11-59 pak2Δ::hphMX ura4+ leu1+ ade6+ his+ | Cross JCS0237 x JCR544 (a gift from JC Ribas) |
| | YSM4544 | h90 cwg1-1 ura4+ leu1+ ade6+ his+ | Cross JCS0237 x JCR296 (a gift from JC Ribas) |
| Figure 6 | YSM4545 | h90 cwg1-1 pak2Δ::hphMX ura4+ leu1+ ade6+ his+ | Cross JCS0237 x JCR296 (a gift from JC Ribas) |

| | | | |
|---|---|---|---|
| | YSM4546 | h90 cwg1-2 ura4+ leu1+ ade6+ his+ | Cross JCS0237 x JCR156 (a gift from JC Ribas) |
| | YSM4547 | h90 cwg1-2 pak2Δ::hphMX ura4+ leu1+ ade6+ his+ | Cross JCS1165 x JCS1166 |
| | YSM4548 | h90 pbr1-8 ura4+ leu1+ ade6+ his+ | Cross JCS0237 x JCR36 (a gift from JC Ribas) |
| | YSM4549 | h90 pbr1-8 pak2Δ::hphMX ura4+ leu1+ ade6+ his+ | Cross YSM4548 x JCS1168 |
| | YSM4550 | h90 pmk1Δ::kanMX myo52-mScarlet-I:natMX lys3+:pmap3:mTagBFP2:termScAdh1:bleMX ade6+:ppck2:RBDpck2-sfGFP:bsdMX ura4+ leu1+ his+ | Cross JCS0889 x JCS0427. pmk1Δ::kanMX comes from PPG4135 (a gift from Pilar Perez) |
| | YSM4551 | h90 pmk1Δ::kanMX pak2Δ::hphMX myo52-mScarlet-I:natMX lys3+:pmap3:mTagBFP2:termScAdh1:bleMX ade6+:ppck2:RBDpck2-sfGFP:bsdMX ura4+ leu1+ his+ | Cross YSM4550 x JCS0428 pmk1Δ::kanMX comes from PPG4135 (a gift from Pilar Perez) |
| | YSM4552 | h90 pmk1Δ::kanMX orb2-34 pak2Δ::hphMX myo52-mScarlet-I:natMX lys3+:pmap3:mTagBFP2:termScAdh1:bleMX ade6+:ppck2:RBDpck2-sfGFP:bsdMX ura4+ leu1+ his+ | Cross YSM4550 x JCS0548 pmk1Δ::kanMX comes from PPG4135 (a gift from Pilar Perez) |
| Figure 7 | YSM4479 | h+ his5+:ptdh1:SynZip4-GFP1-10:natMX ura4+ leu1+ ade6+ | YSM1371 transformed with StuI linearized pSM2952 |
| | YSM4480 | h- myo52-mScarlet-I:natMX ade6+:pact1:SynZip3-GFP11:bsdMX ura4+ leu1+ his+ | Cross JCS0037 x JCS0071 |
| | YSM4481 | h- orb2-34 pak2Δ::hphMX myo52-mScarlet-I:natMX ade6+:pact1:SynZip3-GFP11:bsdMX ura4+ leu1+ his+ | Cross JCS111 x JCS131 |
| Figure S2 | YSM4553 | h90 rho2Δ::kanMX myo52-mScarlet-I:natMX lys3+:pmap3:mTagBFP2:termScAdh1:bleMX ade6+:ppck2:RBDpck2-sfGFP:bsdMX ura4+ leu1+ his+ | Cross YKT013 x JCS0428 rho2Δ::kanMX from bioneer collection |
| | YSM4554 | h90 rho3Δ::kanMX myo52-mScarlet-I:natMX lys3+:pmap3:mTagBFP2:termScAdh1:bleMX ade6+:ppck2:RBDpck2-sfGFP:bsdMX ura4+ leu1+ his+ | osm9328 |
| | YSM4555 | h90 rho4Δ::kanMX myo52-mScarlet-I:natMX lys3+:pmap3:mTagBFP2:termScAdh1:bleMX ade6+:ppck2:RBDpck2-sfGFP:bsdMX ura4+ leu1+ his+ | YSM4457 was transformed with a PCR cassette from plasmid pSM644 and osm9331 osm9332 as primers |
| | YSM4556 | h90 rho5Δ::kanMX myo52-mScarlet-I:natMX lys3+:pmap3:mTagBFP2:termScAdh1:bleMX ade6+:ppck2:RBDpck2-sfGFP:bsdMX ura4+ leu1+ his+ | osm9330 |
| | YSM4557 | h- orb2-34 ura4+ ade6+ leu1+ his+ | Cross YSM4464 x JCS0120 |
