## Supplementary material for "PAK-family kinases promote cell fusion irreversibility by preventing cell wall repair": Table S2

**Table S2: Plasmids used in this study**

| Plasmid N° | Construct | Source |
|---|---|---|
| pSM4231 | pFA6a-3'UTRpak2-5'UTRpak2-pak2-sfGFP-pFA6alpha-KanMX | This study |
| pAV0220 | pFA6a-3'UTRpak2-5'UTRpak2-pFA6a-hphMX | (1) |
| pSM1538 | pFA6a-sfGFP-kanMX6 | (7) |
| pAV0761 | pLys3BstZ17I-pmap3-mTagBFP2-terminatorScADH1-bleMX | (72) |
| pSM2951 | pAde6RsrII/BlpI pact1-SynZip3-GFP11-termScADH1-bsdMX | This study |
| pSM2952 | pHis5StuI-ptdh1-SynZip4-GFP1-10-natMX | This study |
| pSM3006 | pFA6a-3'UTRpak2-5'UTRpak2-pak2ΔPH-CRIB (206-589)-pFA6alpha-kanMX6 | This study |
| pSM3023 | pFA6a-3'UTRpak2-5'UTRpak2-pak2ΔPH-CRIB-KD (K343A, 206-589)-pFA6alpha-kanMX6 | This study |
| pSM3028 | pFA6a-3'UTRpak2-5'UTRpak2-pak2KD (K343A)-sfGFP-pFA6alpha-KanMX | This study |
| pSM3042 | pFA6a-3'UTRpak2-5'UTRpak2-pak2ΔPH (129-589)-sfGFP-pFA6alpha-KanMX | This study |
| pSM3043 | pFA6a-3'UTRpak2-5'UTRpak2-pak2ΔPH-CRIB (206-589)-sfGFP-pFA6alpha-KanMX | This study |
| pSM3044 | pFA6a-3'UTRpak2-5'UTRpak2-pak2ΔKIN (1-279)-sfGFP-pFA6alpha-KanMX | This study |
| pSM3049 | pFA6a-3'UTRpak2-5'UTRpak2-pak2ΔPH-CRIB-K343A (K343A, 206-589)-sfGFP-pFA6alpha-KanMX | This study |
| pSM3057 | pFA6a-3'UTRpak2-5'UTRpak2-pak2HHLL (H137L, H140L)-sfGFP-pFA6alpha-KanMX | This study |
| pSM3067 | pFA6a-3'UTRpak2-5'UTRpak2-pak2x3KA (K33A, K45A, K58A)-sfGFP-pFA6alpha-KanMX | This study |
| pSM3097 | pFA6a-3'UTRpak2-5'UTRpak2-pak2x3KA-HHLL-KD (K33A, K45A, K58A, H137L, H140L, K343A)-sfGFP-pFA6alpha-KanMX | This study |
| pSM3166 | pFA6a-3'UTRpak2-5'UTRpak2-pak2x3KA-HHLL-ΔKIN (K33A, K45A, K58A, H137L, H140L, 1-279)-sfGFP-pFA6alpha-KanMX | This study |
| pSM3191 | pAde6RsrII/BlpI ppck2-pck2RBD(1-381)-sfGFP-termScADH1-bsdMX | This study |
| pSM644 | pFA6a-kanMX6 | (70) |
