## Supplementary material for "PAK-family kinases promote cell fusion irreversibility by preventing cell wall repair": Table S3

### Table S3: Primers used in this study

| osm | sequence |
|---|---|
| 7807 | atgaatcgcaaatcaagtgaatctttggctgacagtcaggattattcgagaaagatattgcgtgtcacaaatttgaaccggatccccgggttaattaac |
| 7808 | tatgaaagtcataaacaccataaagaagaaataattttttggttgaatttaaataaagtaaaaaagaaaaatgttgttgaattcgagctcgtttaaac |
| 8027 | acacctgtcaatactgttctcacacgacaacaacaagagtgtttcagaggattttccagttttgctactgaacggatccccgggttaattaa |
| 8028 | catcagaactatatgaaattgttaccataattttctaattttaaaaaagcagaagcaaaggacaagctaattgaattcgagctcgtttaaac |
| 8031 | gagatgcagcaacattttgaaggtttttcttattcatgcgaggatgataaaccctcgactaccgataatgctcggatccccgggttaattaa |
| 8032 | gaaattagaataatttatcaatgcaatgaaagattaagaaaatgagagtaactttatgctcaatttaaggtggaattcgagctcgtttaaac |
| 8035 | aaaaagcgccatgatcattcttataatgaaactgctgctatagaccataagtctgatgataatcgccataaccggatccccgggttaattaa |
| 8036 | aaaacgcaatgaatgatttgtatgcaaatgttaccgtaaaaaggctcaataaaaccagctgctgccaatgtagaattcgagctcgtttaaac |
| 11580 | ctgacaagaaccatcatttgttc |
| 8030 | gatctctgcaaagataaccc |
| 9327 | attgttacctgaaatgtagaagcc |
| 9328 | gaaacggacagaatcccc |
| 9331 | cgttgtcatggcaatgtttagcttggaaataccagcaatctcagctggtaagcaaacattaacaacaacaaaggcaatcggatccccgggttaattaa |
| 9332 | tgctatacaaataggcgaagcaaaaatgaagacatcaaaaatggtaagattaagtaaataaaggccatgatcaagatagaattcgagctcgtttaaac |
| 9329 | attactctaagacgccgc |
| 9330 | ttaaacattcatggaaactcattagg |
